# Midcell localization of PBP4 of *Escherichia coli* modulates the timing of divisome assembly

**DOI:** 10.1101/2020.07.30.230052

**Authors:** Jolanda Verheul, Adam Lodge, Hamish C.L. Yau, Xiaolong Liu, Xinwei Liu, Alexandra S. Solovyova, Athanasios Typas, Manuel Banzhaf, Waldemar Vollmer, Tanneke den Blaauwen

**Author notes:** contributed equally (Jolanda Verheul, Adam Lodge, Hamish C.L. Yau, Xiaolong Liu).

## Abstract

Insertion of new material into the *Escherichia coli* peptidoglycan (PG) sacculus between the cytoplasmic membrane and the outer membrane requires a well-organized balance between synthetic and hydrolytic activities to maintain cell shape and avoid lysis. The hydrolytic enzymes outnumber the enzymes that insert new PG by far and very little is known about their specific function. Here we show that the DD-carboxy/endopeptidase PBP4 localizes in a PBP1A/LpoA and FtsEX dependent fashion at midcell during septal PG synthesis. Midcell localization of PBP4 requires its non-catalytic domain 3 of unknown function, but not the activity of PBP4 or FtsE. Microscale thermophoresis with isolated proteins shows that domain 3 is needed for the interaction with NlpI, but not PBP1A or LpoA. *In vivo* crosslinking experiments confirm the interaction of PBP4 with PBP1A and LpoA. We propose that PBP4 functions together with the amidases AmiA and B to create denuded glycan strands to attract the initiator of septal PG synthesis, FtsN. Consistent with this model, we found that the divisome assembly at midcell was significantly affected in cells lacking PBP4.

**IMPORTANCE:** Peptidoglycan biosynthesis is a major target for antibacterials. The covalently closed peptidoglycan mesh, called sacculus, protects the bacterium from lysis due to its turgor. Sacculus growth is facilitated by the balanced activities of synthases and hydrolases, and disturbing this balance leads to cell lysis and bacterial death. Because of the large number and possible redundant functions of peptidoglycan hydrolases, it has been difficult to decipher their individual functions. In this paper we show that the DD-endopeptidase PBP4 localizes at midcell during septal peptidoglycan synthesis in *Escherichia coli* and is important for the timing of the assembly of the division machinery. This shows that inhibition of certain hydrolases could weaken the cells and might enhance antibiotic action.

## INTRODUCTION

The peptidoglycan (PG) layer is sandwiched between the cytoplasmic membrane (CM) and the outer membrane (OM) of the Gram-negative bacterium *Escherichia coli* forming a covalently closed network of glycan strands that are interconnected by short peptides[1]. The PG layer maintains the shape of the bacterium by stabilizing the cell against its high internal osmotic pressure. To proliferate, the rod-shaped bacterium grows in length and then divides by binary fission into two equally sized daughter cells. The balanced activities of PG synthases and hydrolases ensures a safe extension of the PG layer without defects that would cause cell lysis[2]. Beta-lactam antibiotics disturb this balance by inactivating the transpeptidase (TPase) activity of the penicillin binding proteins (PBP)s causing lysis of growing cells. We currently know more than twenty PG hydrolases that can cleave either in the glycan chains or peptides within the PG layer, but how their activities are regulated is poorly understood. This is particularly true for the PG endopeptidases[2]. Recently, the OM bound lipoprotein NlpI was shown to interact with similar affinity with several endopeptidases, suggesting that NlpI acts as an interaction hub for endopeptidases to regulate their activities by competing protein-protein interactions.[3].

The basic precursor (lipid II) for PG synthesis is the β-(1,4) linked disaccharide *N*-acetylglucosamine-*N*-acetylmuramic acid (Glc*N*Ac-Mur*N*Ac) with an undecaprenol-pyrophosphate linked to C1 of Mur*N*Ac and a stem peptide with the sequence L-alanine-D-*iso*-glutamate-diaminopimelic acid (Dap)-D-alanine-D-alanine coupled to the C3 of Mur*N*Ac. Lipid II is used by glycosyltransferases (GTases), which polymerize the glycan chains, and TPase, which crosslink the stem peptides. The main PG synthases in *E. coli* are the class A PBPs (PBP1A and PBP1B) with GTase and TPase activities [4–8], the integral membrane proteins RodA and FtsW with GTase activity[9–11] and their partner TPases PBP2 and PBP3, respectively[12–15].

The hydrolases are present in greater redundancy than the synthases[2,16]. The carboxypeptidases (CPase) PBP5, PBP6A and PBP6B remove the terminal D-Ala from nascent pentapeptides[17-20]. The endopeptidases (EPases) PBP4, PBP7, MepA, MepM, MepH, and MepS cleave the crosslinks in the PG[21-23]. The amidases AmiA-D hydrolyze the bond between Mur*N*Ac and L-alanine at position 1 of the stem peptide[24-27]. Finally, the lytic transglycosylases Slt70 and MltA-G cleave the glycan strands to release 1,6-anhydroMur*N*Ac containing turnover products[28-30]. The large number of hydrolases reflects the adaptability to various environmental conditions {Pazos; Muellersw} but begs the question of how their potentially dangerous activities are controlled to avoid cell lysis.

Binary fission in *E. coli* is initiated by the assembly of the Z-ring at midcell, which consist of a network of dynamic FtsZ filaments that are anchored to the cytoplasmic membrane by FtsA and ZipA[31-32]. The Z-ring also recruits the PG synthases PBP1A and PBP1B to midcell[4-33]. A number of Z-associated proteins play a role in the organization and dynamics of the ring[34-39]. Of these, the integral membrane protein FtsX interacts with FtsA[40] and its cytoplasmic ATPase partner protein FtsE interacts with FtsZ[41]. Both localize early to midcell, together with the Z-ring[24]. As a complex FtsEX recruits EnvC, which activates AmiA and AmiB that are needed for septum cleavage[26,42]. FtsE versions without ATPase activity still localize at midcell and recruit as FtsEX-EnvC complex but the latter is not able to activate the amidases[43]. Presumably an ATPase dependent conformational change in FtsEX induces a conformational change in EnvC, which is then able to relieve an autoinhibitory alpha-helix from the active site of AmiA and AmiB, activating them[44]. The third amidase AmiC is activated by the OM-anchored lipoprotein NlpD[24] and all three amidases AmiA-C are activated under certain stress conditions by the recently identified ActS[46,47]. The recruitment of the other cell division proteins occurs with a time delay of about 20% of the cell division cycle, depending on the growth condition[48-50]. FtsK, FtsBLQ, FtsW, PBP3 and FtsN localize in an interdependent fashion. FtsBLQ inhibit the PG synthase activity of PBP3, FtsW and PBP1B until they are outcompeted by the accumulation of FtsN that relieves this inhibition and initiates septum synthesis[51-54].

While investigating the specific function of various PG hydrolases, we found that PBP4 localizes specifically at midcell as part of the division machinery. PBP4 is a periplasmic endopeptidase[55] with a C-terminal amphipathic alpha-helix that associates with membranes[56]. Deletion of *dacB*, which encodes PBP4, does not have morphological consequences but causes a decrease in the percentage of PG crosslinks and of PG-attached lipoprotein[57] and an increased sensitivity to bile salts[58]. Overproduction of PBP4 is toxic[59] presumably because the enhanced cleavage of peptide cross-links weakens the PG mesh and eventually causes lysis. PBP4 has three domains that are assembled in an unusual way[60] (Fig. 1A). A non-catalytical domain of unknown function (domain 2) is inserted into the transpeptidase domain 1, and a third domain (domain 3) is inserted into domain 2. Domain 3 is positioned above the active site with its catalytic serine 62, which resides in domain 1, and might be involved in substrate binding or regulation[61] (Fig. 1A). Site directed mutagenesis suggested that residues from domain 1 (R361) and 2 (D155) are important for the endopeptidase activity[62].

**Fig 1.**
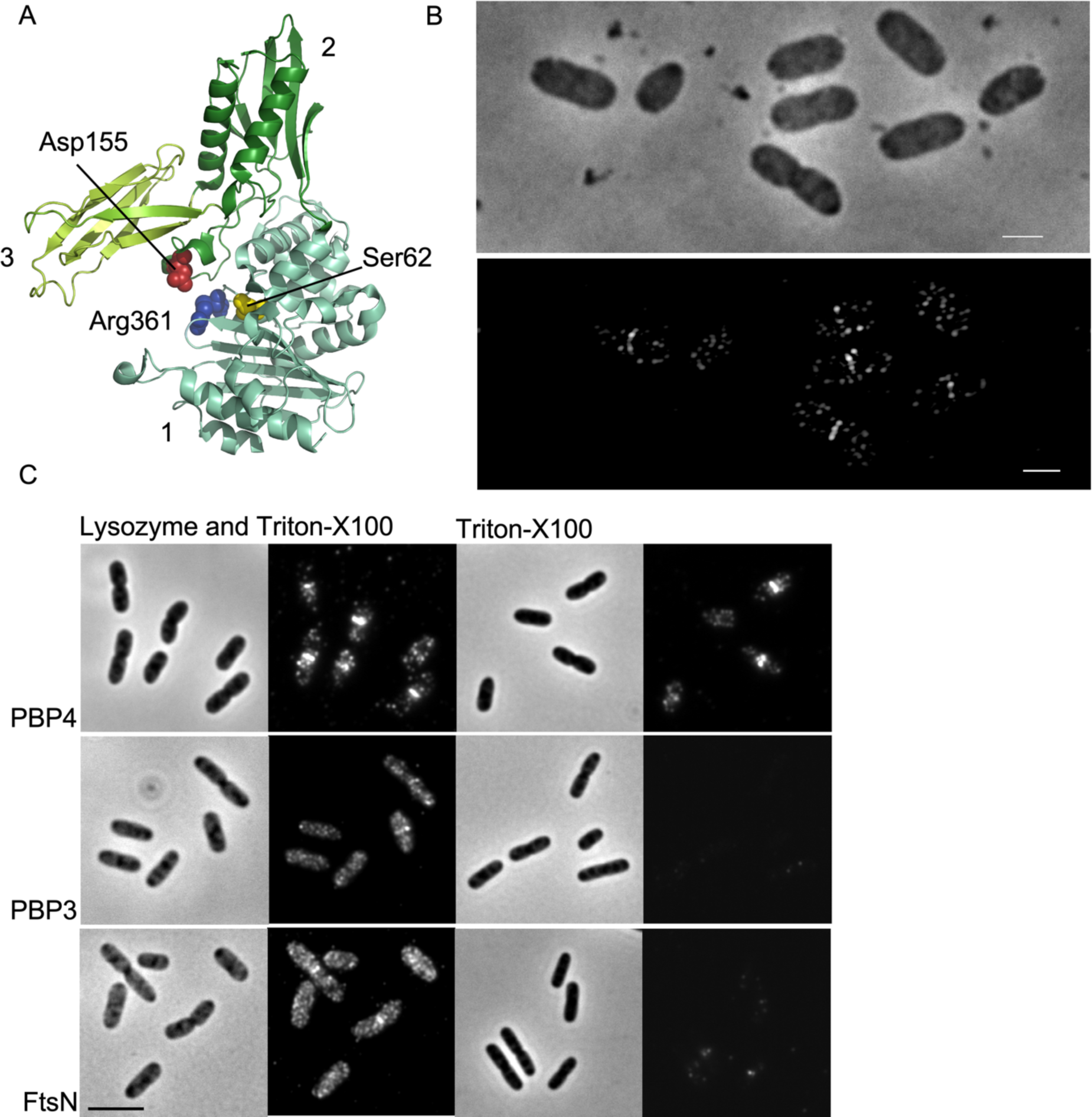
Structure of PBP4 and its localization at midcell and in the lateral wall during fast growth. A. Crystal structure of PBP4 in which the three domains (numbers 1-3) and some of the residues that are essential for its endopeptidase activity are indicated. B. MC4100 cells were grown exponentially in TY medium at 37°C to an OD_600_ of 0.3 and fixed and immunolabeled with BW25113Δ*dacB* pre-adsorpted anti-PBP4. Upper panel phase contrast image corresponding to the SIM fluorescence image in the lower panel. Scale bar equals 2 µm C. BW25113 wild-type cells were grown in TY at 37°C and harvested in the exponential phase at an OD_600_ of 0.3 and fixed. Samples were divided into three aliquots and cells were immunolabeled with antibodies against PBP3, FtsN and PBP4. The first aliquot was immunolabeled without any treatment, the second after permeabilizing the cells with TritonX-100 (right: phase contrast and fluorescence images) and the third after permeabilizing cells with Triton-X100 and lysozyme (left: phase contrast and fluorescence images). The scale bar equals 5 µm. PBP4, but not PBP3 or FtsN, is accessible without degrading the peptidoglycan layer.

Here we determined the localization of PBP4 and different variants during the cell cycle of *E. coli.* The timing of localization at midcell and interactions of PBP4 support a role in remodeling of PG synthesized by the divisome.

## RESULTS

### PBP4 localizes to the lateral wall and at midcell

To determine the localization of PBP4 in *E. coli* cells as a function of the bacterial cell cycle, we generated an antiserum against PBP4 and removed unspecific antibodies by adsorption to cells of a *dacB* deletion mutant that lacks PBP4. The purified antibodies were specific for PBP4 (Fig. S1) and used to immunolabel the wild-type strain MC4100 grown in rich (TY) medium at 37°C. PBP4 localized strongly at midcell and to a lesser extend in the lateral wall in wild-type cells (Fig. 1B).

### PBP4 localizes between the peptidoglycan layer and the outer membrane

Although it was reported that PBP4 has a stretch of weakly amphipathic amino acids at its C-terminus[56], it was not known whether PBP4 freely diffuses in the periplasm or associates with the membrane or other proteins. It has been previously observed that chemically fixing freely diffusing periplasmic proteins can shock them into the cell poles[12-63]. The fact that we did not observe any polar localization of PBP4 after fixation (Fig. 1B and S1), suggests that it is either associated with the membrane or other proteins. Next, we investigated whether PBP4 was accessible to anti-PBP4 in cells with an intact PG layer. For this, wild-type cells were grown in TY at 37°C and immunolabeled without the typically used Triton X-100-mediated permeabilization of the OM or without digesting the PG mesh with lysozyme. PBP4 was not accessible without permeabilization of the OM but was fully accessible in cells with an intact PG (Fig. 1C). In contrast, proteins like FtsN and PBP3 were not accessible without degradation of the PG layer by lysozyme (Fig. 1C). The OM pore inserting BAM complex is also not accessible without lysozyme treatment[64]. This suggests that PBP4 resides between the OM and the PG layer.

### Activity of PBP4 is not needed for midcell localization

Substrate binding can be a key determinant in the localization of proteins [18-65-66]. To test whether PBP4 substrate binding is needed to localize at division sites we expressed the active site mutants PBP4(S62G) and (S62A)[61] that fails to bind β-lactams and therefore is not able to hydrolyze the PG stem peptide (Fig. S2 and S3). A plasmid encoding the mutant protein was constitutively expressed in a Δ*dacB* background by the leakage of a weakened p*Trc99*A promoter. Interestingly, the wild-type and active site mutant when expressed from plasmid were both able to localize at midcell (Fig. 2). This indicates that although PBP4(S62G) cannot hydrolyze its substrate or bind β-lactams covalently (Fig. S2C Fig. S3), it is either still able to interact with its substrate or it localizes through protein interactions.

**Fig 2.**
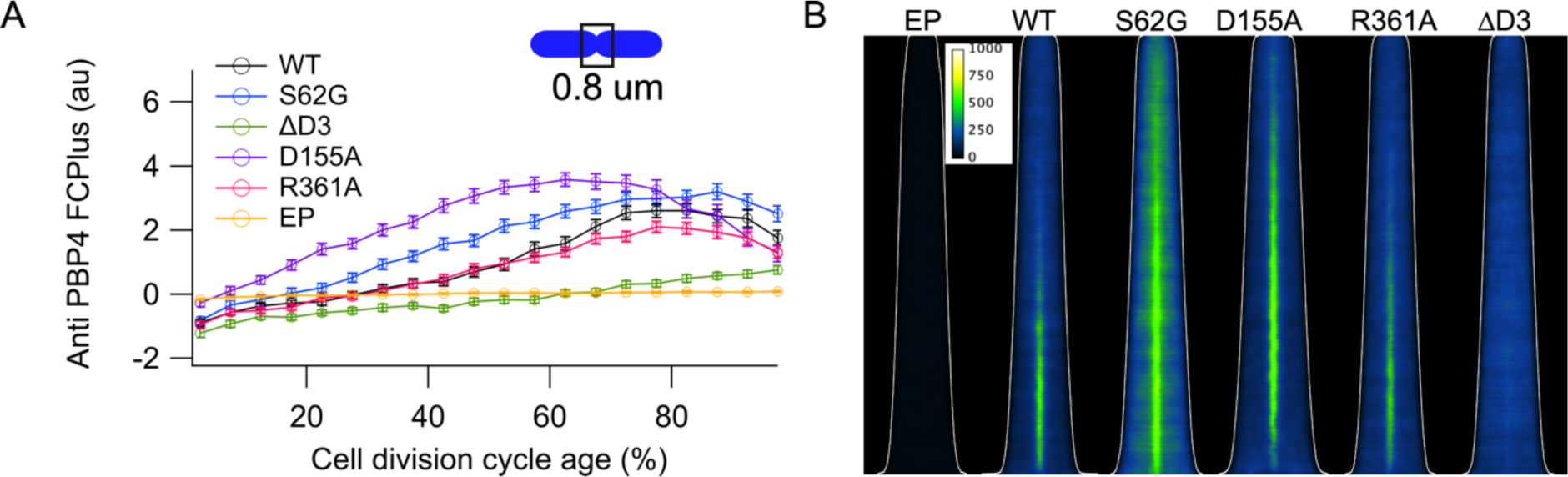
Activity of PBP4 is not required for localization. Cells were grown in minimal glucose medium to steady state at 28°C, fixed and immunolabeled with antibodies against PBP4. (A) The extra fluorescence at midcell (FCPlus) in the Δ*dacB* strain transformed with the empty plasmid (EP, yellow), or plasmids expressing wild-type PBP4 (black), PBP4S63G (blue), PBP4D155A, (purple) PBP4R361A (red), PBP4ΔD3 (green) was determined and plotted as function of the cell division cycle age as in bins of 5% age classes with the error bar indicating the 95% confidence interval. (B) Demographs of the localization fluorescence pattern of the PBP4 variants shown in (A) with the cells sorted according to their cell length. The white line indicates the length of the cells. Intensity scaling is identical for all maps. Number of cells analyzed for each immunolabeling was at least 2000 cells.

Since PBP4(S62G) could still interact with its PG substrate, we mutagenized the gene to express PBP4 versions in which residues D155 or R361 involved in the binding of the DAP residue in the stem peptide by alanine[62] were replaced. The genes encoding PBP4(D155A) or PBP4(R361A) were expressed by promoter leakage from plasmid in a Δ*dacB* background, and both PBP4 versions localized at midcell (Fig. 2). An immunoblot analysis showed that all mutants were expressed to similar levels and that PBP4(R361A) and PBP4(D155A) were able to bind the fluorescent β-lactam Bocilin-FL (Fig. S2). Therefore, all three active site protein variants are potentially able to interact with substrate as an inactive protein.

A PBP4 version lacking domain 3 (PBP4ΔD3) was still able to bind β-lactams (Fig. S2B and S3) but did not have DD-carboxypeptidase or DD-endopeptidase activity (Fig. S3). Analytical ultracentrifugation (Fig. S4A) and circular dichroism analysis (Fig. S4B) of isolated PBP4ΔD3, showed that the removal of domain 3 did not affect the structure, the dimerization of the rest of the protein or its interaction with PG sacculi (Fig. S4C), which suggests a folded active site domain. However, although the inactive PBP4ΔD3 was capable of binding substrate, it did not localize at midcell (Fig. 2). Hence, we conclude that the midcell localization of PBP4 is largely driven by protein-protein interactions and to a lesser extent by substrate interaction but is independent from its own activity.

### PBP4 localizes at inactive divisomes

To further dissect whether the localization of PBP4 at the division site is dependent on the availability of its substrate or the presence of other proteins at the division site, cell division was inhibited through the specific inactivation of PBP3 by aztreonam[67]. Aztreonam treatment for 1-3 mass doublings stalls the division machinery at midcell resulting in filamentous growth. In the longer filaments new division machineries are formed and localize at future division sites[5]. In these filaments PBP4 still localized albeit somewhat weaker than in untreated cells at all possible division sites (Fig. S5), indicating that the activity of PBP3 was not required for the recruitment of PBP4. While PBP3 is inhibited, preseptal PG synthesis[68] still occurs at the potential division sites in the filamenting cells [18,33,69]. The pre-septal localization of PBP4 mimics the localization of PBP1A and PBP1B, which were previously shown to be recruited to preseptal PG synthesis sites by early cell division proteins[4,33].

### PBP1A/LpoA assist in midcell localization of PBP4

PBP4 was recently reported to interact with PBP1A, LpoA and the EPase adaptor protein NlpI[3]. To investigate whether PBP1A and/or LpoA proteins were needed for midcell localization of PBP4, strains deleted for *mrcA* (PBP1A), *lpoA*, *mrcB* (PBP1B) or *lpoB* were grown in rich medium at 37°C and immunolabeled with anti-PBP4. The amount of PBP4 that localized at midcell was identical for the parental wild-type strain BW25113, Δ*mrcB* and Δ*lpoB,* but reduced in the Δ*mrcA* and Δ*lpoA* strains (even after correction for the smaller diameter of these cells,[70], whereas the PBP4 concentration was similar for all strains (Fig. 3). This could indicate that PBP1A/LpoA assist in the initial localization of PBP4 at midcell.

**Fig 3.**
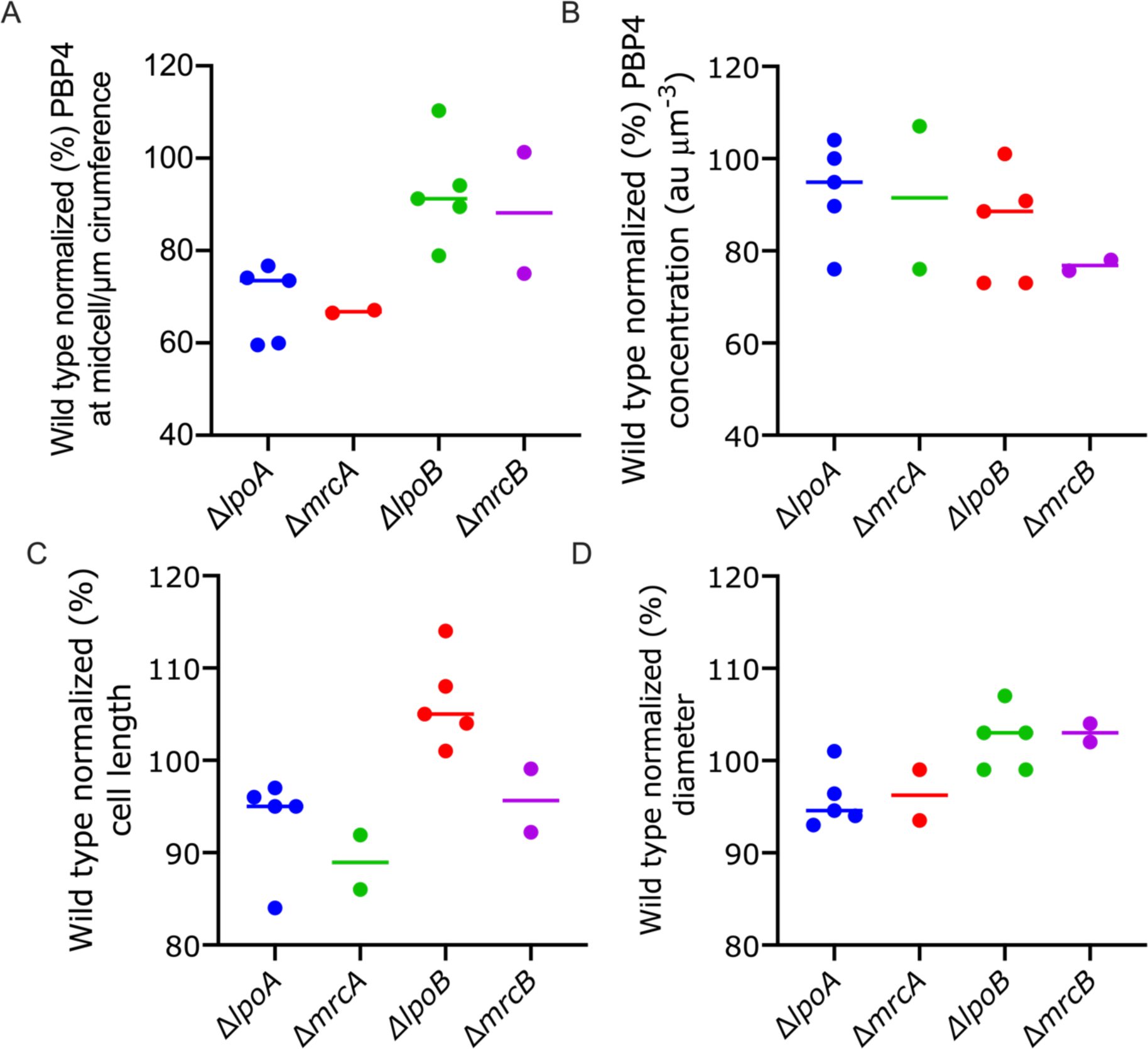
PBP1A and LpoA contribute to PBP4 midcell localization. Cells were grown exponentially to an OD of 0.3 in TY at 37°C, then fixed and immunolabeled with antibodies against PBP4. Because fluorescence imaging by microscopes is usually not directly comparable between different experiments, all results in each experiment were normalized to the parental strain BW25113. (A) PBP4 fluorescence at midcell per μm circumference of the cell. (B) Concentration of PBP4 in the cells. (C) length of the cells. (D) Diameter of the cells. Δ*lpoA* (n=5), Δ*lpoB* (n=5), Δ*mrcA* (PBP1A, n=2), Δ*mrcB* (PBP1B, n=2). Each point is the average of 1000-2000 cells. Based on the one-way Anova the difference in midcell localization is significant (P=0.009) while the difference in concentration of PBP4 is not significant (P=0.0484).

### NlpI seems not to be involved in midcell localization of PBP4

We next tested if the interaction with the OM anchored NlpI affected the localization of PBP4. In the Δ*nlpI* cells the amount of PBP4 at midcell per μm average cell circumference was reduced by 42 ± 6.7% (n = 3) of the wild type and the protein localized later at midcell than in the wild-type cells (Fig. S6). However, the PBP4 concentration in the Δ*nlpI* strain was also lower than in the wild-type cells (64.8 ± 17.6%, n = 3). This could indicate that the cells down-regulate PBP4 expression or that PBP4 becomes unstable in the Δ*nlpI* strain. Labeling of PBP4 in a Δ*nlpI* strain did not require lysozyme (Fig. S6), suggesting that NlpI is not needed to maintain PBP4 localized outside the PG layer, which could be facilitated by additional interactions of PBP4.

### PBP4 domain 3 facilitates interaction with NlpI but not PBP1A/LpoA

Microscale thermophoresis (MST) in which one protein is labelled with a fluorophore and titrated with an unlabeled partner protein was used to analyze the interactions between PBP4 and its known interaction partners further. The difference in thermophoresis of the complex versus the unbound fluorescently labeled proteins is then used to calculate the affinity of the binding partners. Different versions of PBP4 and its interaction partners were purified and assayed for direct protein-protein interactions. Wild-type PBP4 interacted with PBP1A, LpoA and NlpI[3]. PBP4 lacking domain 3 (PBP4ΔD3), which did not localize at midcell (Fig. 2), still interacted with PBP1A and LpoA but not with NlpI (Fig. 4). These results suggest that domain 3 of PBP4 facilitates the interaction with NlpI, but not PBP1A.

**Fig 4.**
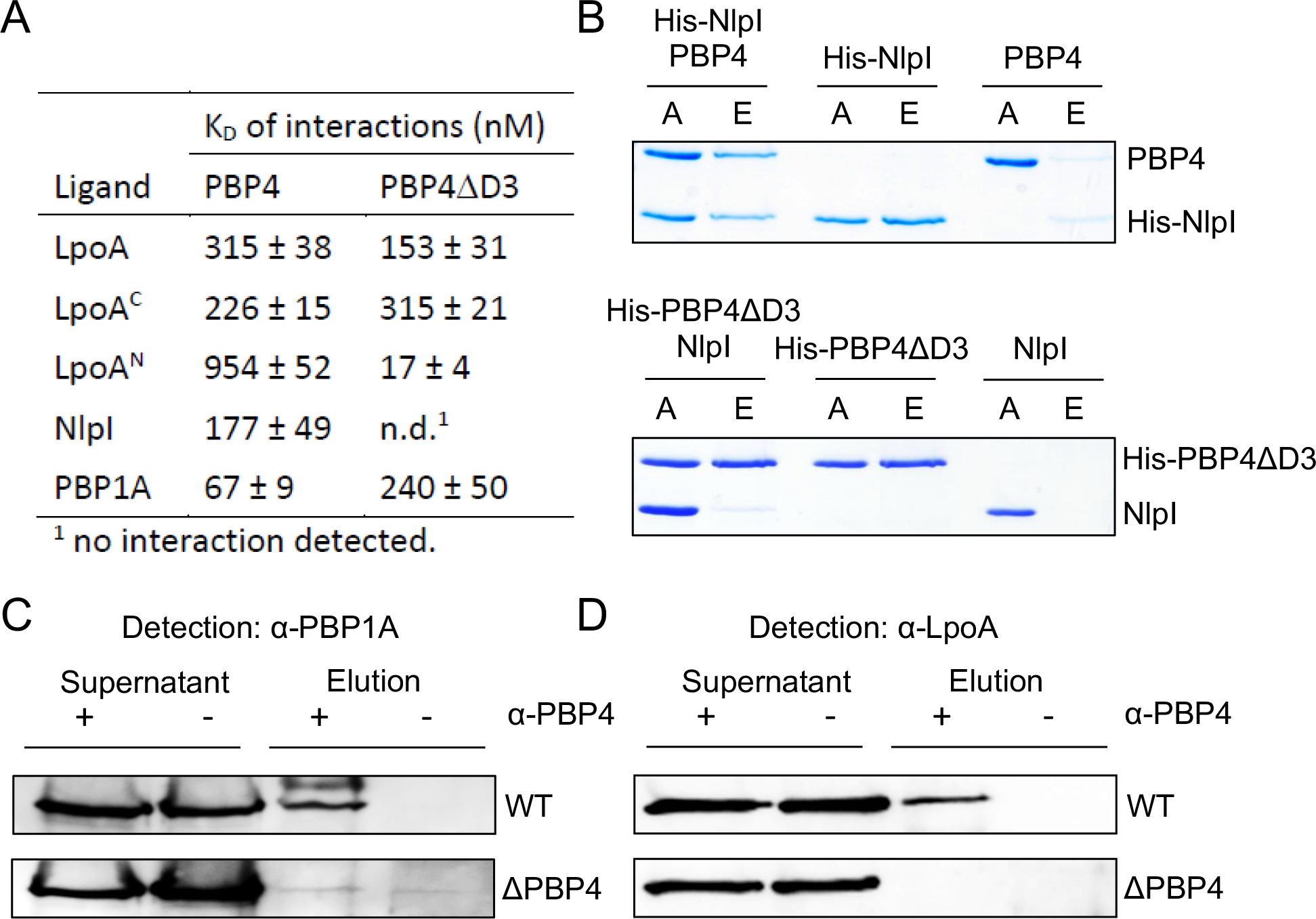
PBP4 interacts with PBP1A, LpoA and NlpI. (A) Summary of apparent K_D_ values of the interactions of PBP4 or a version lacking domain 3 (PBP4ΔD3) with LpoA, the C-terminal domain of LpoA (LpoA^C^), the N-terminal domain of LpoA (LpoA^N^), NlpI and PBP1A determined by microscale thermophoresis (MST). Corresponding binding curves are shown in Suppl. Figure S7. (B) Pulldown experiment showing that domain 3 of PBP4 is required for interaction with NlpI. Oligo-histidine tagged NlpI (His-NlpI) pulled down PBP4 to Ni-NTA beads. His-tagged PBP3ΔD3 did not pull down untagged NlpI. A, applied sample; E, eluted sample. (C) *In vivo* cross-linking/co-immunoprecipitation showing interaction between PBP1A and PBP4. Growing cells of BW25113 (wt) or BW25113Δ*dacB* (ΔPBP4) were chemically cross-linked with DTSSP and cell extract was immunoprecipitated with purified anti-PBP4 antibody. Control samples did not receive anti-PBP4 antibody. The cross-linker was cleaved by reducing agent and proteins were separated by SDS-PAGE and transferred to a membrane, followed by detection of PBP1A with specific antibodies. PBP1A was detected in the elution of the sample from wild-type and not from ΔPBP4. (D) *In vivo* cross-linking/co-immunoprecipitation (as in panel C) showing that PBP4 and be cross-linked with LpoA in cells.

### NlpI and PBP1A/LpoA have different binding sites in PBP4

PBP4 can form a multi-enzyme complex with NlpI and PBP1A/LpoA[3]. PBP1A interacted with PBP4 and could be cross-linked to PBP4 in cells (Fig. 4C). We found that full length LpoA interacted with both PBP4 and PBP4ΔD3, with apparent K_D_ values of 315 ± 38 nM and 153 ± 31 nM, respectively (Fig. 4A; Fig. S7). This suggests that domains 1 or 2 of PBP4 are sufficient for the interaction with LpoA. MST assays of PBP4 or PBP4ΔD3 with LpoA’s C-terminal (residues N257-S678; LpoA^C^) or N-terminal (LpoA residues G28-T256; LpoA^N^) domains also yielded positive binding curves (Fig. 4A; Fig. S7)[71-73]. We found that LpoA^C^ and the full length LpoA version had comparable affinities for both PBP4 versions, suggesting that the C-terminal domain of LpoA is sufficient for interaction with PBP4 (Fig. 4A). Interestingly, the removal of domain 3 from PBP4 resulted in an increased affinity of LpoA^N^ for PBP4 (app. K_D_ decreased by ∼50 fold) (Fig. 4A Fig. S7). Hence, the interaction between the N-terminal domain of LpoA and PBP4 possibly occurs via a different mechanism dependent on a conformational rearrangement of domain 3 of PBP4. This suggests that the two domains of LpoA, which likely serve different primary functions, are both sufficient for an interaction with PBP4, albeit with varying affinities. In conclusion PBP4 robustly interacts with PBP1A/LpoA independent of its domain 3.

### LpoA has a small effect on PBP4 activity

We found previously that NlpI does not affect PBP4 activity[3] but considered it possible that PBP4 interaction with PBP1A/LpoA may affect its activity. We first verified that NlpI or PBP4 did not affect the activity of PBP1A in the presence or absence of LpoA using an *in vitro* glycosyltransferase assay (Fig. 5). We next tested if PBP1A or LpoA affected the activity of PBP4 using *in vitro* PG degradation assays. LpoA decreased the activity of PBP4 by about 30% in a continuous DD-CPase assay with soluble substrate and an endpoint PG digestion assay (Fig 5B and C). To assess whether LpoA also affects PBP4 activity *in vivo*, a Δ*dacB*Δ*lpoA* strain and Δ*dacB* strain were transformed with the plasmids expressing PBP4 or the active site mutant PBP4(S62G) and the toxicity of their expression was compared. Although the double mutant was less fit than the single mutant, it was equally sensitive to the overexpression of PBP4 (Fig. 5D), leaving no indication that LpoA also inhibits the activity of PBP4 *in vivo*.

**Fig 5.**
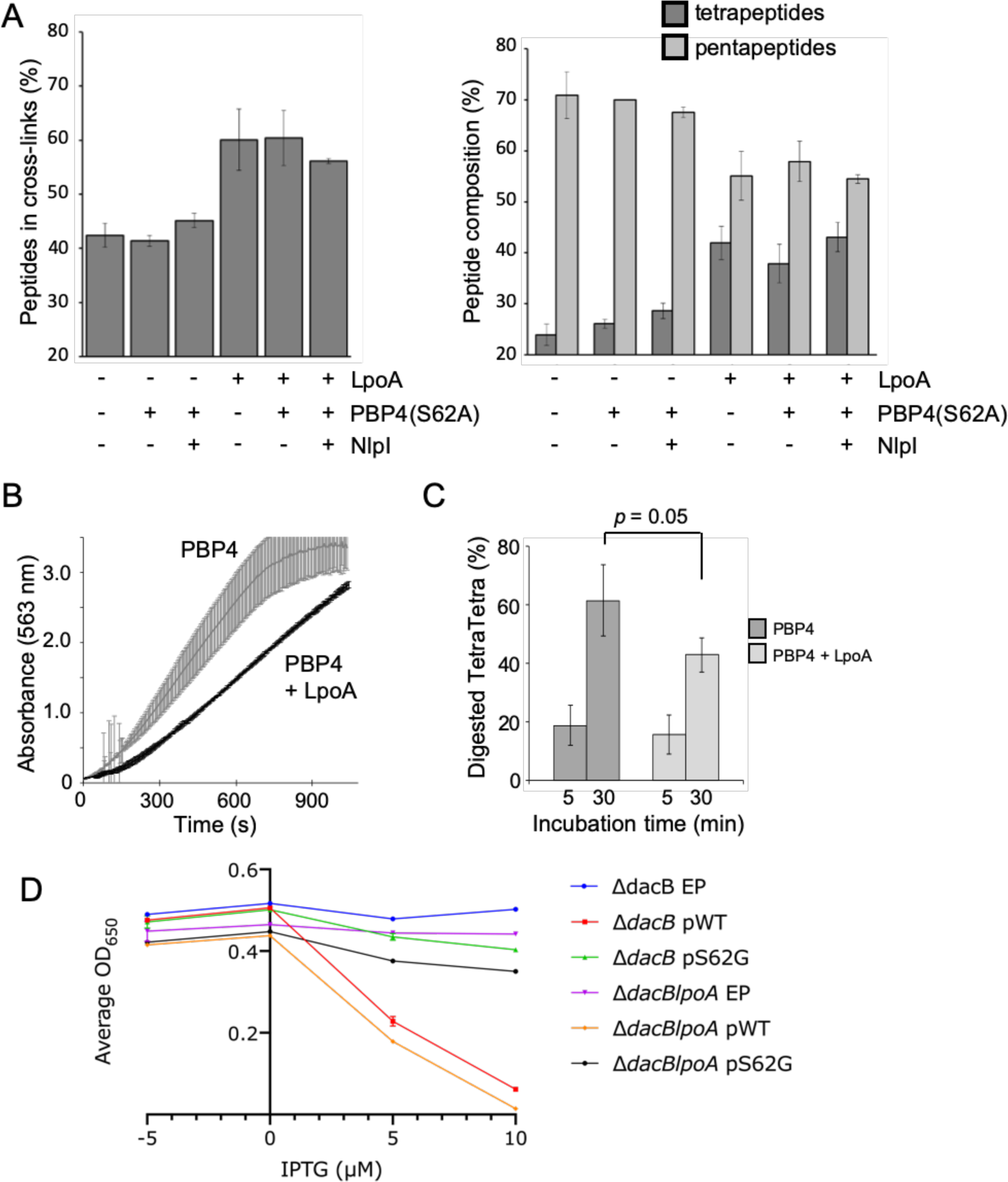
PBP4 or NlpI do not affect the activity of PBP1A whereas LpoA reduces the activity of PBP4. (A) Summary of the results from an *in vitro* PG synthesis assay with PBP1A (in the presence or absence of LpoA) and radio-labelled lipid II substrate. The presence of NlpI, catalytically inactive PBP4(S62A) or both together, does not affect the cross-linkage of the PG product of a PBP1A or PBP1A/LpoA reaction (left side), and NlpI, PBP4(S62A) or both together, don’t induce enhanced carboxypeptidase activity of PBP1A, which would be seen as a higher content of tetrapeptides (right side). Values are mean ± variation of two independent repeats. Example chromatograms are shown in Figure S8. (B) The presence of LpoA modestly reduces the activity of PBP4 in a DD-carboxypeptidase assay using the substrate *N*-acetyl-L-Lys-D-Ala-D-Ala. (C) LpoA also reduces slightly the activity of PBP4 against PG sacculi from BW25113, as seen by the reduced digestion of TetraTetra upon a 30 min incubation period. (D The Δ*dacB* and Δ*dacB*Δ*lpoA* mutant strains are equally sensitive to overexpression of PBP4. The average growth over time as measured by OD650 nm is plotted against the added IPTG concentration. The culture grown in TY with glucose to suppress the expression is represented by -5 on the X-axe. All other data points are from cell grown in TY without glucose (n=4).

### PBP4 localizes as early divisome protein

Since PBP4 localizes at midcell it could be part of or associated with the divisome. The division machinery is assembled in two successive steps[48,74]. First, FtsZ and its membrane bound partners ZipA and FtsA in conjunction with other Z-ring associated proteins form the proto-ring that localizes at midcell {denBlaauwen:2017dq}. Second, after approximately 20% of the cell division cycle the remaining proteins assemble subsequently at the division site. To obtain more details on PBP4 localization in relation to other cell division proteins, MC4100 cells were grown to steady state in minimal glucose medium at 28°C. The cells were immunolabeled with antibodies against PBP4, FtsZ (early localizing), or FtsN) late localizing). PBP4 localized strongly at midcell and hardly in the lateral cell wall (Fig. 6). To determine whether PBP4 belongs to the early or late divisome localizing proteins, we analyzed the timing of its midcell localization. To this end, we plotted the extra fluorescence (FCPlus) present at midcell in comparison to the amount of fluorescence in the rest of the cell (Fig. 6). PBP4 localization coincided with that of FtsZ, indicating that PBP4 localizes relatively early.

**Fig 6.**
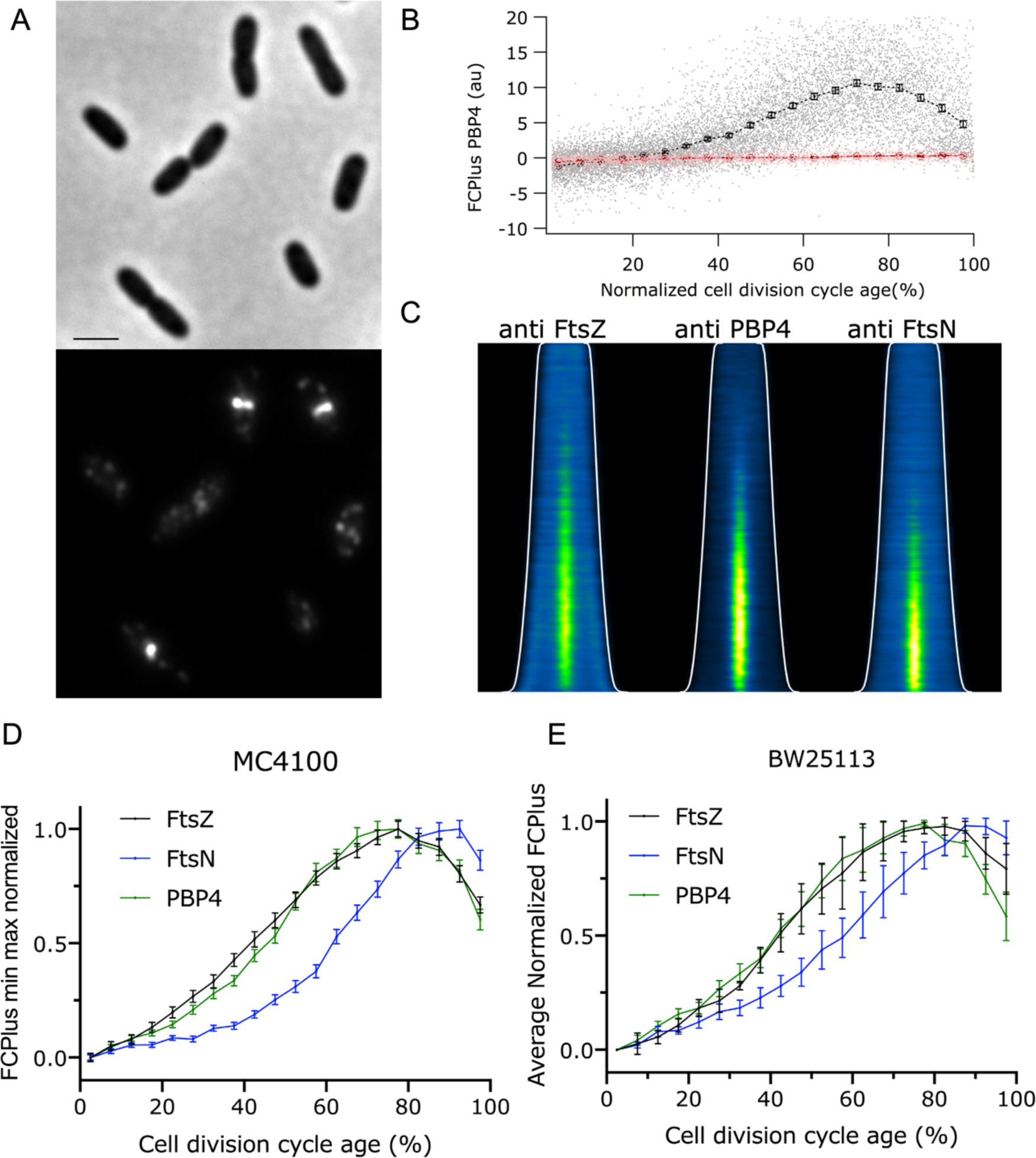
Cell division cycle timing of the localization of PBP4. (A) MC4100 cells were grown to steady state in minimal glucose medium at 28°C and immunolabeled with antibodies specific for PBP4. Phase contrast (top) and fluorescence is shown (bottom). Scale bar equals 2 µm. (B) The extra fluorescence at midcell compared to the rest of the cell (FCPlus) is plotted as function of the normalized cell division cycle age. The dots (grey for MC4100 and light pink for the BW25113 Δ*dacB strain*) are the values measured for the individual cells and the markers with bars (black for the wild-type MC4100, red for the BW25113 Δ*dacB*) are 5% age bins with 95% confidence. (C) Demographs of the localization of FtsZ, PBP4 and FtsN in the cells sorted according to cell length. The white line indicates the length of the cells. (D) Comparison of the timing of the localization at midcell of FtsZ (black), PBP4 (green) and FtsN (blue) during the cell division cycle age. The FCPlus values are min-max normalized to enable timescale comparison, despite differences in molecule number and antibody affinities. (E) BW25113 wild type cells were grown to steady state in minimal glucose medium at 28°C and immunolabeled with antibodies specific for FtsZ, PBP4 and FtsN. The FCPlus of three independent biological experiments in 5% age bins was determined. The binned FCPlus values were min-max normalized. The average min-max normalized FCPlus values of the three experiments were subsequently plotted as function of the cell division cycle.

### Midcell localization of PBP4 depends on FtsE

To investigate if PBP4 localization is dependent on the presence of the Z-ring or the later localizing division associated proteins we localized PBP4 in various division defective strains. First, the localization of PBP4 was assessed in a strain that expressed Tre1 that ADP-ribosylates FtsZ at R174, which renders it unable to participate in protofilament formation[75,76]. The inactive E415Q variant of Tre1 was used as negative control. PBP4 was unable to localize in cells with insufficient FtsZ protofilaments to form division Z-rings (Fig. 7A), whereas it localized as in WT cells in the presence of the TreE415Q variant, proving that a Z-ring is needed for PBP4 localization. Next, strains harboring temperature sensitive alleles of FtsE, FtsQ and PBP3 (FtsI) were grown in minimal glucose medium at 28°C to an OD_450_ of 0.2, diluted 1:4 in prewarmed medium of 28°C or 42°C and grown for two mass doublings before PBP4 immunolabelling. PBP4 localized in filamentous cells with thermo-labile FtsQ or PBP3 indicating that these proteins are not needed to recruit PBP4 (Fig. 7B). However, PBP4 localized very diffuse around potential division sites in the filamentous *ftsE*(ts) cells (Fig. 7B). This suggests that a functional FtsE is important for PBP4 localization. The FtsEX/EnvC complex assists in the assembly of the divisome[77,78] and is essential for amidase function to cleave PG for cell separation[43]. To investigate a possible link between PBP4 and this complex we further dissected the localization requirement of PBP4. PBP4 localized at division sites in a Δ*envC* strain, indicating that EnvC is not needed for PBP4 localization. However, PBP4 failed to localize in a Δ*ftsEX* strain (Fig. 8). To verify whether FtsE and FtsX were required for PBP4 localization we constructed a Δ*ftsE* strain. The *ftsE* and *fts*X genes reside in an operon and the expression of *fts*X requires the *fts*E gene[79]. We therefore constructed a Δ*ftsE* strain with a weakened p*Trc*99A promoter placed upstream of *fts*X to ensure sufficient expression of FtsX. The resulting strain, XL36 grew with a mild filamentous phenotype in rich medium that could be complemented by the expression of FtsE(wt) (Fig S9). PBP4 did not localized at septal positions in the Δ*fts*E strain XL36 (Fig. 8A) but localized normally when FtsE was expressed to complement this strain (Fig 8B). This suggests that FtsE and FtsX enhance the septal localization of PBP4. Interestingly, PBP4 localized at division sites in a strain lacking the three amidases AmiA, AmiB and AmiC (Fig. S10) and in a strain lacking essential components of the Tol-Pal system, which is involved in the coordination of OM constriction and PG invagination during cell division (Fig. S10).

**Fig 7.**
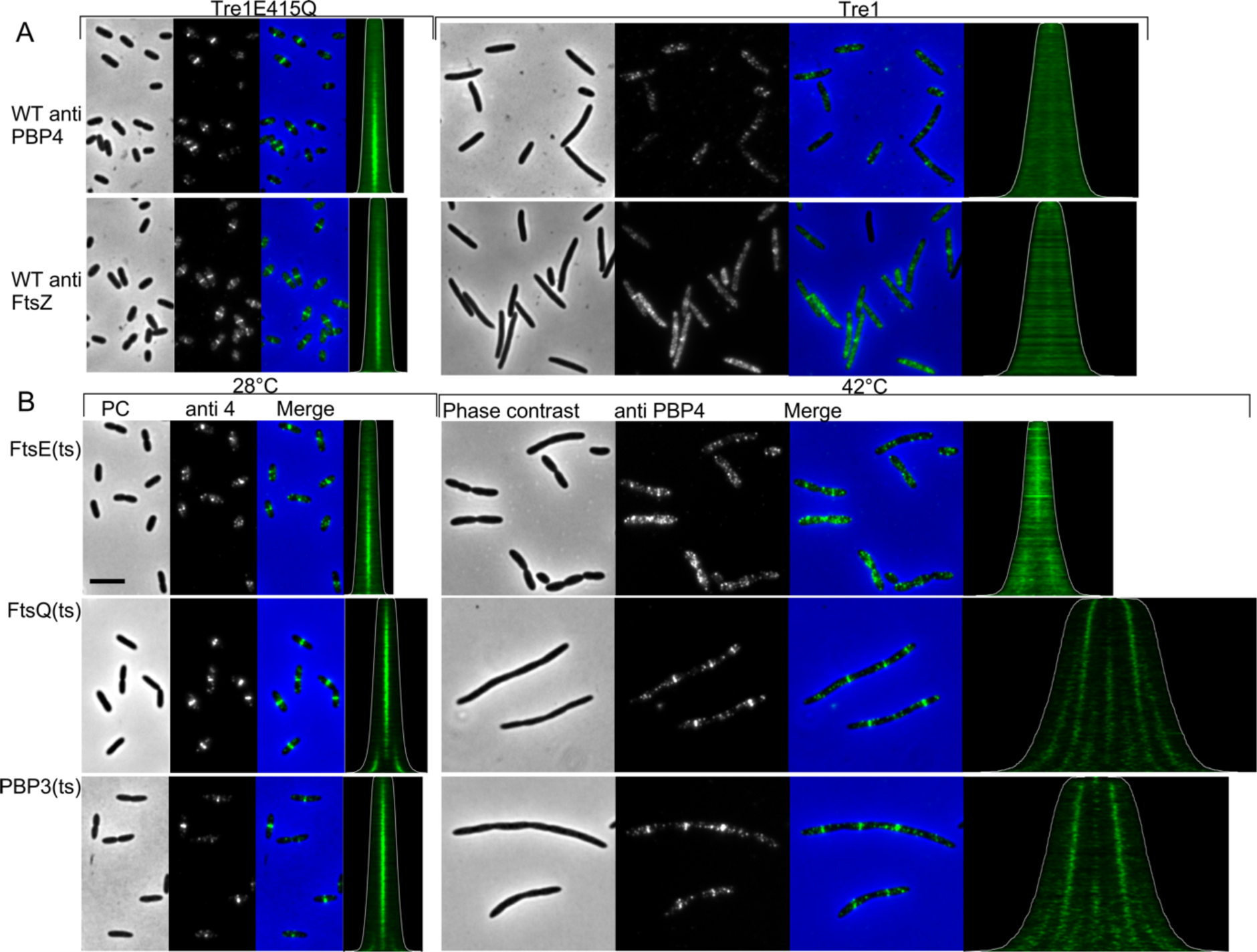
PBP4 localization is dependent on the presence of the proto-ring. **A.** MC4100 was transform with a plasmid that expressed either Tre1 that abolished the ability of FtsZ to polymerize by ADP-ribosylation of residue R174 or with the inactive variant Tre1(E415Q). Cells were grown in Gb1 medium at 28°C and expression of the inhibitor was induced for 2 mass doublings (MD) with 0.15% arabinose. Subsequently, the cells were fixed and immunolabeled with antibodies against FtsZ or PBP4. From left to right: the phase contrast (PC), corresponding fluorescence image of the PBP4 labeling and the merged images ate shown. In the demographs the cells are sorted according to cell length (contours in white). The number of cells analyzed were 3694 and 3756 for Tre1E415Q and 1275 and1621 for Tre1 for anti FtsZ and anti PBP4, respectively. **B.** Isogenic strains of MC4100 producing different temperature sensitive (ts) versions of cell division proteins were grown to steady state in Gb1 medium at 28°C and split in two parts. These were 1:4 diluted in prewarned medium and grown for two MD at either 28°C or 42°C. The cells were fixed, labeled with antibodies specific for PBP4 (a4). From left to right: the phase contrast (PC), corresponding fluorescence image of the PBP4 labeling and the merged images at the permissive temperature, demograph of PBP4 fluorescence localization where cells are sorted according to their cell length (contours in white). This is followed by the same series from the non-permissive temperature samples. The number of cells analyzed were 2809 and 1576 for LMC515 FtsE(ts), 3833 and 1203 for LMC531 FtsQ (ts) and 3876 and 926 for LMC510 PBP3(ts), for the cells grown at 28°C and 42°C, respectively. The scale bar equals 5 µm. Within one antibody staining the brightness and contrast of the samples is identical and therefore directly comparable.

**Fig 8.**
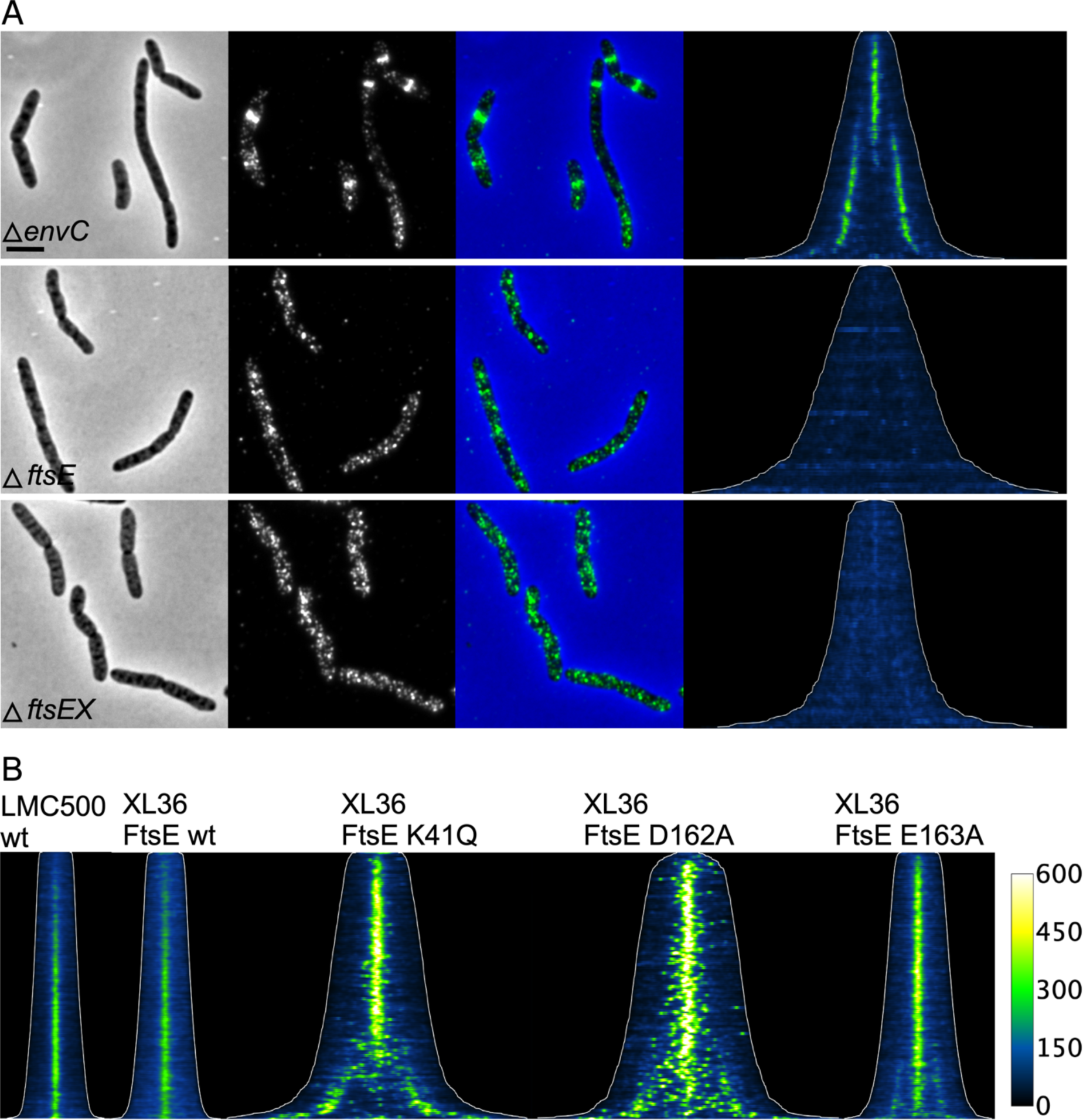
PBP4 is recruited by the septal cleavage complex FtsEX. (A) Cells of BW25113 expressing the proteins indicated were grown in LB at 37°C to OD_600_ = 0.3, fixed and labeled with specific antibodies against PBP4. From left to right, the phase contrast, corresponding fluorescence image of the PBP4 labeling, and the merged former two images, map of fluorescence with the cell lengths outlined (with the same contrast and brightness for all maps) of PBP4 localization where cells are sorted according to their cell length are shown. The number of cells analyzed were 687 for Δ*envC*, 434 for Δ*ftsE*, and 346 for Δ*ftsEX*. The scale bar equals 5 μm. (B) Cells of LMC500 or XL36 (LMC500Δ*ftsE*::p*Trc99A*down *fts*X) expressing wild-type or active site versions of FtsE were immunolabeled with anti-PBP4 antibodies. Cells were grown in TY at 30°C and FtsE expression from plasmid was induced for 2 mass doubling with 30 μM IPTG, and cell were fixed and harvested at an OD_600_ of 0.3. The maps of PBP4 fluorescence with the cell lengths outlined is shown for the indicated strains. Number of cells analyzed: LMC500 (2462), (wt) (2588), K41Q (728), D162A (554), and E163E (1052).

To determine whether the activity of FtsE was needed for the localization of PBP4, we expressed three inactive FtsE variants that had been shown to localize to midcell {Arends:2009id}. The K41Q and D162A versions of FtsE cannot bind ATP and FtsE(E163A) binds ATP but cannot hydrolyze it. PBP4 was able to localize at the division site in cells that expressed one of these inactive variants of FtsE as the only copy (Fig. 8B) showing that the activity of FtsE is not required for the localization of PBP4.

### PBP4 affects the timing of the divisome assembly

To measure the timing of divisome in various strain backgrounds, we grew Δ*dacB* cells carrying either an empty plasmid or plasmids constitutively expressing PBP4 or PBP4 versions to steady state in minimal glucose medium at 28°C. Cells were fixed, divided into three aliquots and labeled with antibodies against FtsZ, PBP4 or FtsN. We determined the timing of the arrival of these proteins at midcell by analyzing fluorescence signals of several thousands of cells[80]. In the parental strain FtsZ and FtsN started to localize at midcell at about 26 % and 49%, respectively, of the cell cycle age. Similarly, in the Δ*dacB* strain expressing wild-type PBP4 from plasmid FtsZ and FtsN started to localize at midcell at about 28% and 49%, respectively, of the cell division cycle age (Fig. 9E). Expression of inactive PBP4 versions affected the timing of the divisome assembly differently. Especially the Δ*dacB* cells expressing PBP4(D155A) from plasmid became slightly filamentous (Fig. 9B), assembled the divisome very early and showed a longer cell division period (Fig. 9E). Next, we measured the timing of the divisome assembly in the absence of PBP4 in the ΔdacB strain and in its parental BW25113. A difference was observed between the two strains, which was not significant due to the large variation in the localization timing of FtsZ and FtsN in both strains (Fig 9F). We inferred that BW25113 is not able to reach a reproducible steady state, possibly due to a known frameshift in the *rph* gene, which leads to pyrimidine starvation and hence irregular DNA replication[81]. The uracil and thymine that we add as standard to the minimal medium for this strain is evidently unable to fully compensate the deficiency. To solve this possible issue, the *dacB* gene was deletion from the wild-type strain MC4100 and the cells were grown to steady state in minimal medium in the absence of an antibiotic. The results reproduced the difference between the Δ*dacB* and wild-type strains but now with a significance of P = 0.0001 (Fig. 9F). No significant difference between the Δ*dacB* and its parental in other parameters that could account for a change in cell cycle such as mass doubling time, cell length, diameter or the percentage of constricting cells was detected (Table S1). However, FtsZ started to localize (t_0_) 8 min earlier in the Δ*dacB* strain and reached 50% of its maximum intensity (t_1/2_) 9 min earlier compared to the parental strain. FtsN localized 8.4 min (t_0_) and 5.7 min (t_1/2_) earlier (see for an explanation of t_0_ and t_1/2_ Fig. 9D). These results show that PBP4 profoundly affects the timing of divisome assembly.

**Fig 9.**
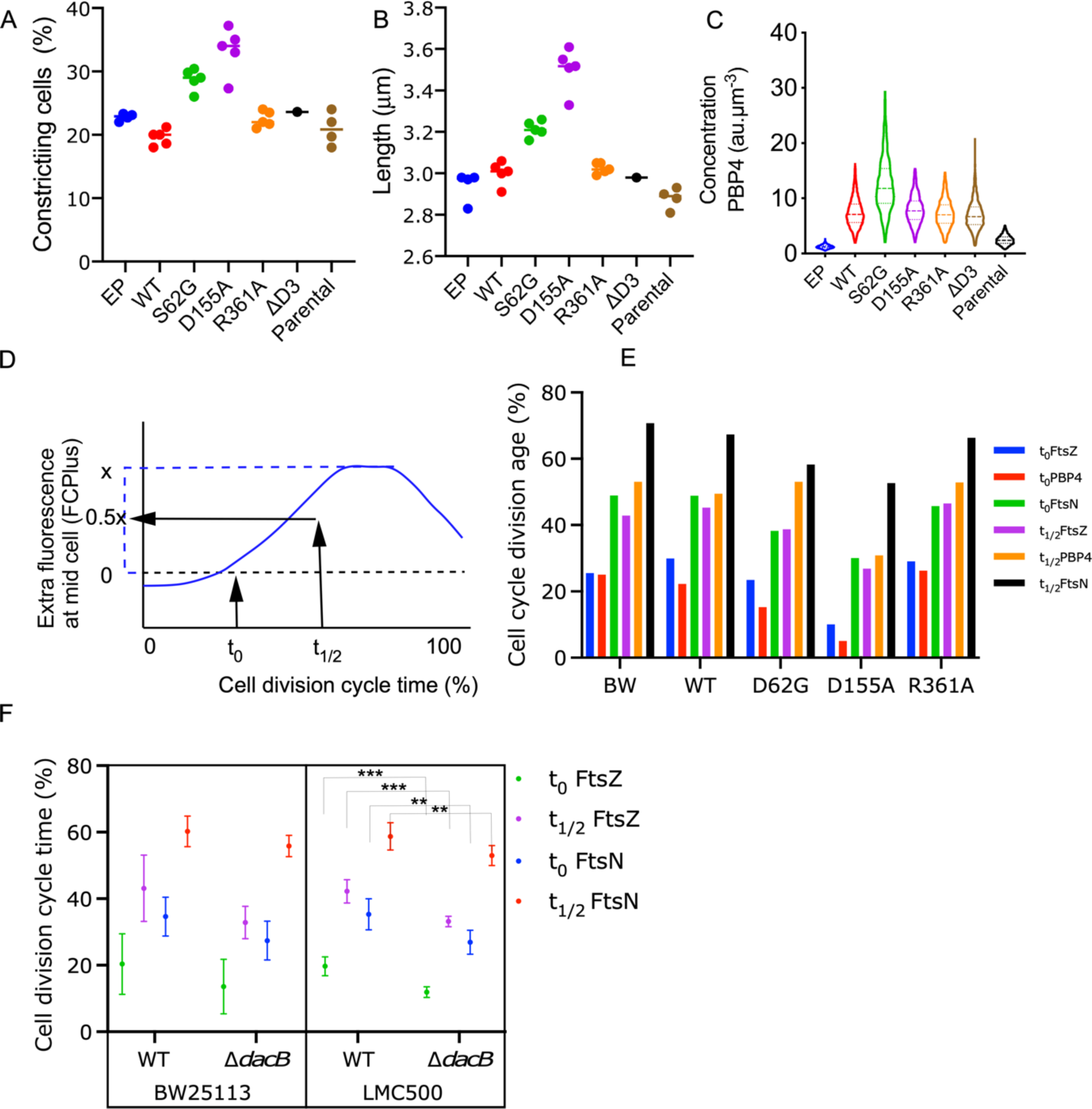
Localization of PBP4 depends on the presence of domain 3 but not on activity. Cells were grown in minimal glucose medium to steady state at 28°C, fixed and immunolabeled with the indicated antibodies. (A and B) Graphs with the percentage of constricting cells and the average cell length in μm for the various mutants expressed from plasmid in the Δ*dacB* strain and also of the parental strain BW24113 (n = 4). (C) The concentration of PBP4 in fluorescent units per μm^3^ in these cells for a representative experiment (out of the four repeats). (D) Graphical illustration of the meaning of t_0_ and t_1/2_. (E) Cell division cycle age timing of FtsZ, PBP4, and FtsN for the parental strain BW25113, the Δ*dacB* strain carrying plasmids for the expression of PBP4WT, PBP4S63G, PBP4D155A, or PBP4R361A. Plasmids were not induced. (F). Cell division cycle age timing of FtsZ and FtsN for the Δ*dacB* strain and its parental BW25113 (left) and the Δ*dacB* strain and its parental LMC500. P 0.0001 = ***, P 0.001 = **. Number of points correspond to the number of independent biological replicates.

## DISCUSSION

*E. coli* contains a large number of seemingly redundant PG hydrolases, which possibly fine tune peptidoglycan biogenesis and remodelling in response to environmental parameters. Apart from their enzymatic activities, little is known about the actual function of most PG hydrolases. Here we report that the DD-endopeptidase PBP4 localizes at midcell during septal PG synthesis in *E. coli* and is important for the timing of the assembly of the division machinery.

### PBP4 localises at midcell during cell division in an FtsEX dependent manner

PBP4 localizes predominantly at midcell during cell division in an FtsEX dependent manner. Inactive PBP4 variants were able to localize at midcell indicating that substrate hydrolysis is not a requirement for localization. However, a version of PBP4 lacking the non-catalytic domain 3 (PBP4ΔD3), which is required for activity but not β-lactam binding, did not localize to mid-cell. From these results we infer that PBP4 localizes to midcell either by recognizing its substrate (without the need to hydrolyse it) and/or by protein-protein interactions. PBP4 molecules appear to arrive early and concomitantly with the assemblage of the proto-ring and may associate to preseptal PG synthesis, which is consistent with our observation that PBP4 affects the timing of divisome assembly.

### PBP4 might be involved in preseptal PG synthesis

PBP4 interacts with NlpI, PBP1A and LpoA and these proteins are able to form a ternary complex ([3] and (Fig. 4). PBP4ΔD3 interacted with PBP1A and LpoA but not with NlpI, confirming differential PBP4 binding sites of PBP1A, LpoA and NlpI. The interaction with PBP1A and/or LpoA may contribute to the recruitment of PBP4 at midcell, as the amount of PBP4 at midcell was slightly reduced in Δ*mrcA* and Δ*lpoA* strains. PBP1A and its partner LpoA are involved in preseptal PG synthesis[33-80]. Although somewhat weaker, PBP4 still localizes at putative division sites in cell filaments generated with aztreonam that have lost septal synthesis but not preseptal synthesis activity. Possibly, some PBP4 enriches at preseptal PG synthesis sites together with PBP1A/LpoA, which localize through the interaction of PBP1A with ZipA[33], and preseptal PG synthesis provides substrates for PBP4. The interaction between NlpI and PBP4 could function to sequester PBP4 in the lateral wall to prevent its interaction with potential substrate (Fig. 10).

**Fig 10.**
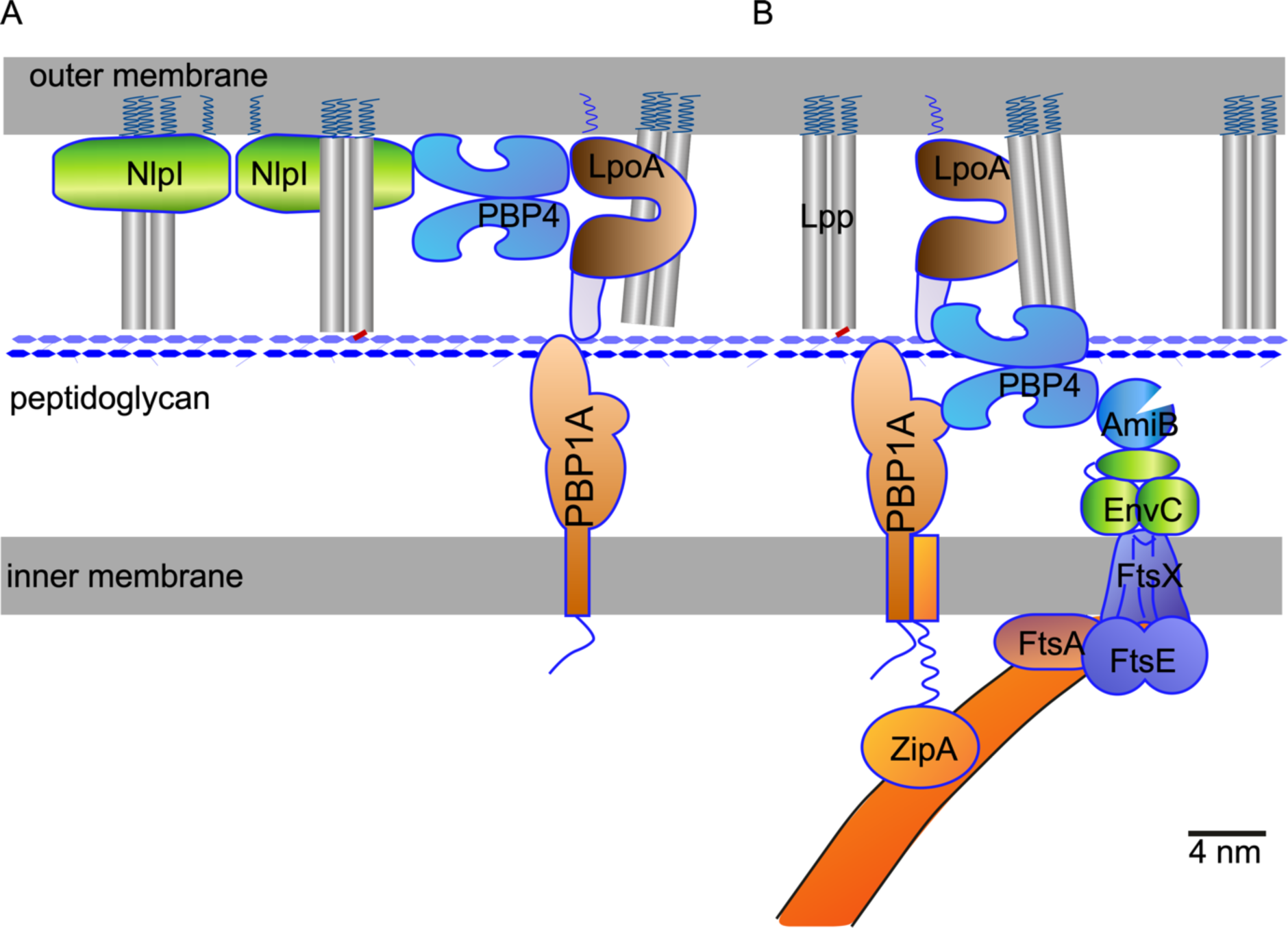
Model of the organization of PBP4 in the periplasm of *E. coli*. In a minimal medium grown cell of average size of 3 by 0.86 μm 230000 Lpp trimeric proteins are present of which 1/3 is bound to PG. (A) During elongation PBP4 (or a fraction of the PBP4 molecules) is kept away from its substrate through its interaction with NlpI and possibly LpoA. (B) PBP1A and LpoA associate with ZipA and the Z-ring to assist in preseptal PG synthesis. Because of their presence, the absence of NlpI and the newly synthesized PG, PBP4 is attracted by PBP1A and LpoA. More PBP4 is accumulating at midcell and becoming clearly visible by immunolabeling upon maturation of the divisome and synthesis of septal PG. All proteins apart from the FtsZ polymer have been drawn according to their crystal structure shape and hydrated crystal structure sizes (scale bar equals 4 nm). The membranes are assumed to be 2 nm and the distance between the outer and the inner membrane 21 nm.

### Why does the loss of PBP4 affects the timing of divisome assembly?

The absence of PBP4 interferes with the timing of the divisome assembly. In most bacteria studied thus far, the Z-ring with its associated early division proteins localizes first to prepare the future cell division site, perhaps by generating the border between the side wall and the future septum [33,68,82]. After a time-delay, the proteins that synthesize the bulk of the septal PG arrive and septation starts[48-50]. In the absence of PBP4, the assembly of the divisome starts earlier in the cell cycle and it takes more time. This change of divisome assembly could be caused by direct or indirect effects. PBP4 could be directly needed to open the PG layer to allow insertion of new septal material or to remodel the nascent PG to allow new insertions. Alternatively, the absence of PBP4 might enable NlpI to bind other EPases, coordinating their activity for remodelling the PG layer during septation. However, the robust localization of PBP4 at midcell suggests a direct function for PBP4 in cleavage of existing glycan crosslinks to allow the insertion of new material. Since deletion of PBP4 does not have dramatic effects, another endopeptidase must be involved as well or be able to replace PBP4 to a large extend. We hypothesise that during elongation sufficient PBP4 molecules are retained by NlpI in the lateral wall above the PG layer to avoid promiscuous PBP4 activity (Fig. 10). This is consistent with the observations that the absence of NlpI resulted in a reduced amount of PBP4 present in the cells (Fig. S6 and[3]). NlpI is not enhanced at midcell[3] and therefore we do not anticipate a role for NlpI in the retention of PBP4 at midcell. PBP4 might be initially recruited to midcell through interactions with PBP1A and LpoA, and the FtsEX-EnvC complex, likely directing PBP4’s activity already to preseptal sites (Fig. 10). Once septal peptidoglycan synthesis starts by the mature divisome, more substrate should become available for PBP4 facilitating its accumulation at midcell, which we observed in the immunolocalization experiments. Hence, PBP4 could initially remove crosslinks to help AmiA and AmiB to generate denuded glycan strands. These would facilitate the accumulation of FtsN, which is required to initiate septal PG synthesis. Alternatively, PBP4 could simply remove old glycan strands during preseptal PG synthesis. Because of its impact on the timing of the divisome assembly, the first option seems to be more likely. Whether the function of PBP4 remains the same during septation will need further investigation.

## MATERIALS AND METHODS

### Culturing conditions

*E. coli* K12 cells were grown to steady state[80] in glucose minimal medium containing 6.33 g of K_2_HPO_4_.3H_2_O, 2.95 g of KH_2_PO_4_, 1.05 g of (NH_4_)_2_, 0.10 g of MgSO_4_.7 H_2_O, 0.28 mg of FeSO_4_.7 H_2_O, 7.1 mg of Ca(NO_3_)_2_·4 H_2_O, 4 mg of thiamine and 4 g of glucose per L. For strain MC4100 and its derivatives 50 µg lysine (Gb1) and for BW25114 and MG1655 derivatives 2 µg uracil, 20 µg thymidine, 50 µg arginine and glutamine (Gb4) per liter pH 7.0 at 28°C or 37°C were added. Absorbance was measured at 450 nm with a 300-T-1 spectrophotometer (Gilford Instrument Laboratories Inc.). Alternatively, cells were grown in TY (5g yeast extract, 10 g bacto trypton and 85 mM NaCl, pH 7.0), or LB, which is TY with 170 mM NaCl and absorbance was measured at 600 nm.

### Strain construction

Strain XL35, the FtsE partial deletion strain, was constructed as follows: Primers priXL221 and priXL222 were used to amplify the upstream homologous sequences (*UHS*) from the *ftsE* start codon, and primers priXL223 and priXL224 were used to amplify the downstream homologous sequence after the FtsE 151^th^ amino acid (the potential promoter of *ftsX*), from the MC4100 genome. Primers priXL51 and priXL54 were used to amplify the kanamycin resistance cassette from plasmid pKD4[83]. The amplified overlap PCR product (*UHS-FRT-KAN-FRT-DHS*) with primers priXL221 and priXL224 was used to construct the recombinant strain XL34 in which the first 150 amino acids encoding sequence of *ftsE* was deleted[83]. Subsequently, the kanamycin selection cassette was removed with plasmid pCP20[83] yielding XL35.

Strain XL36, the *ftsE* clean deletion strain, was constructed similarly as XL35. Primers priXL221 and priXL222 were used to amplify the upstream homologous sequences from the *ftsE* start codon, and primers priXL225 and priXL226 were used to amplify the downstream homologous sequences after the *ftsE* stop codon, from the MC4100 genome. In order to restore the FtsX expression, a *pTrc99Adown* promoter[84] was amplified from plasmid pSAV057[85] with primers priXL77 and priXL227 and a chloramphenicol selection cassette was amplified from plasmid pKD3 with primers priXL52 and priXL54. The amplified overlap PCR product (*UHS-FRT-CAM-FRT-DHS*) with primers priXL221 and priXL226 was used to construct the recombinant strain XL33 in which the entire *ftsE* coding sequence was deleted and FtsX was expressed from the chromosomally encoded p*Trc99Adown* promoter. Subsequently, the kanamycin selection cassette was removed with plasmid pCP20 yielding XL36 (Table 1).

**TABLE 1.**
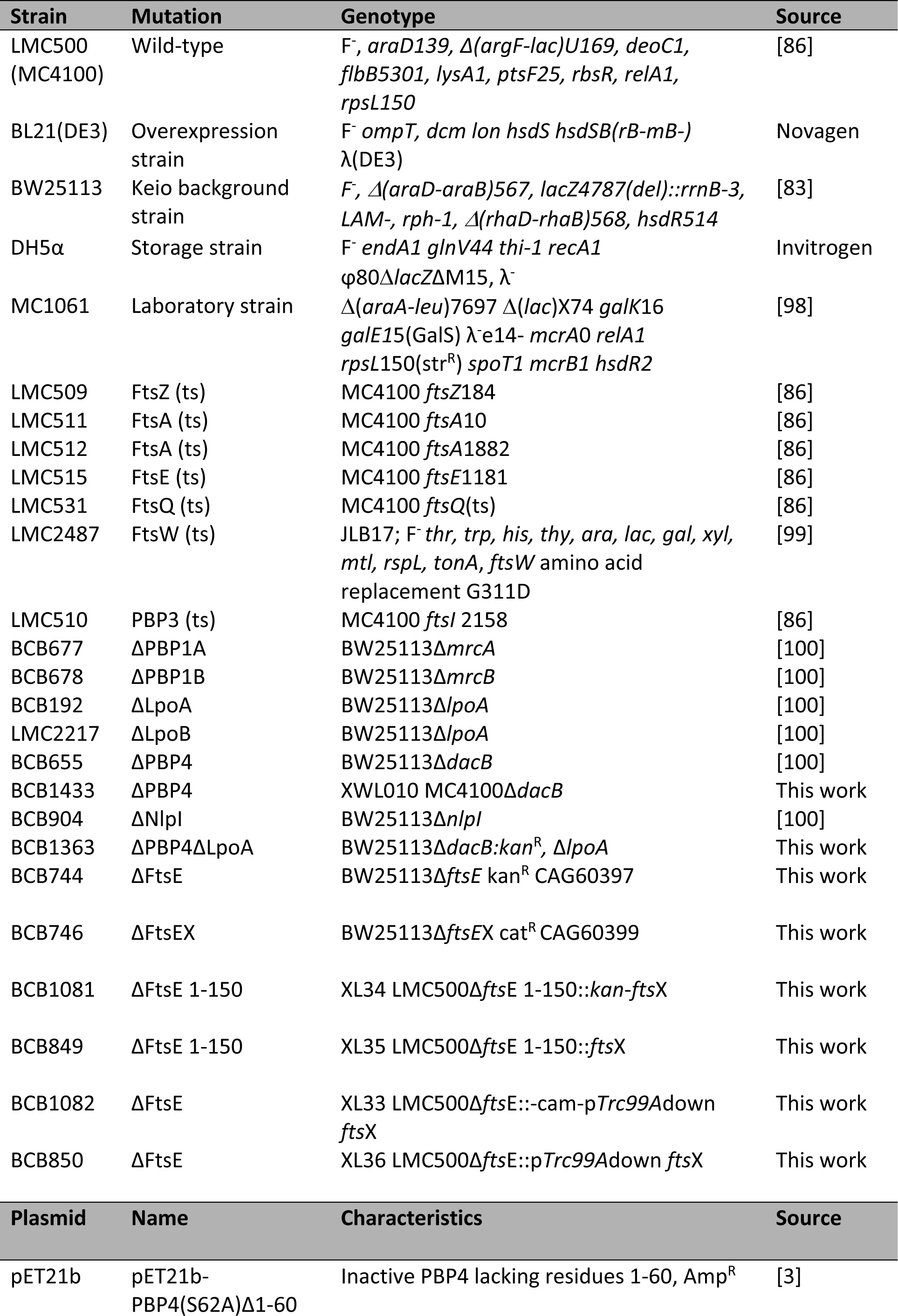
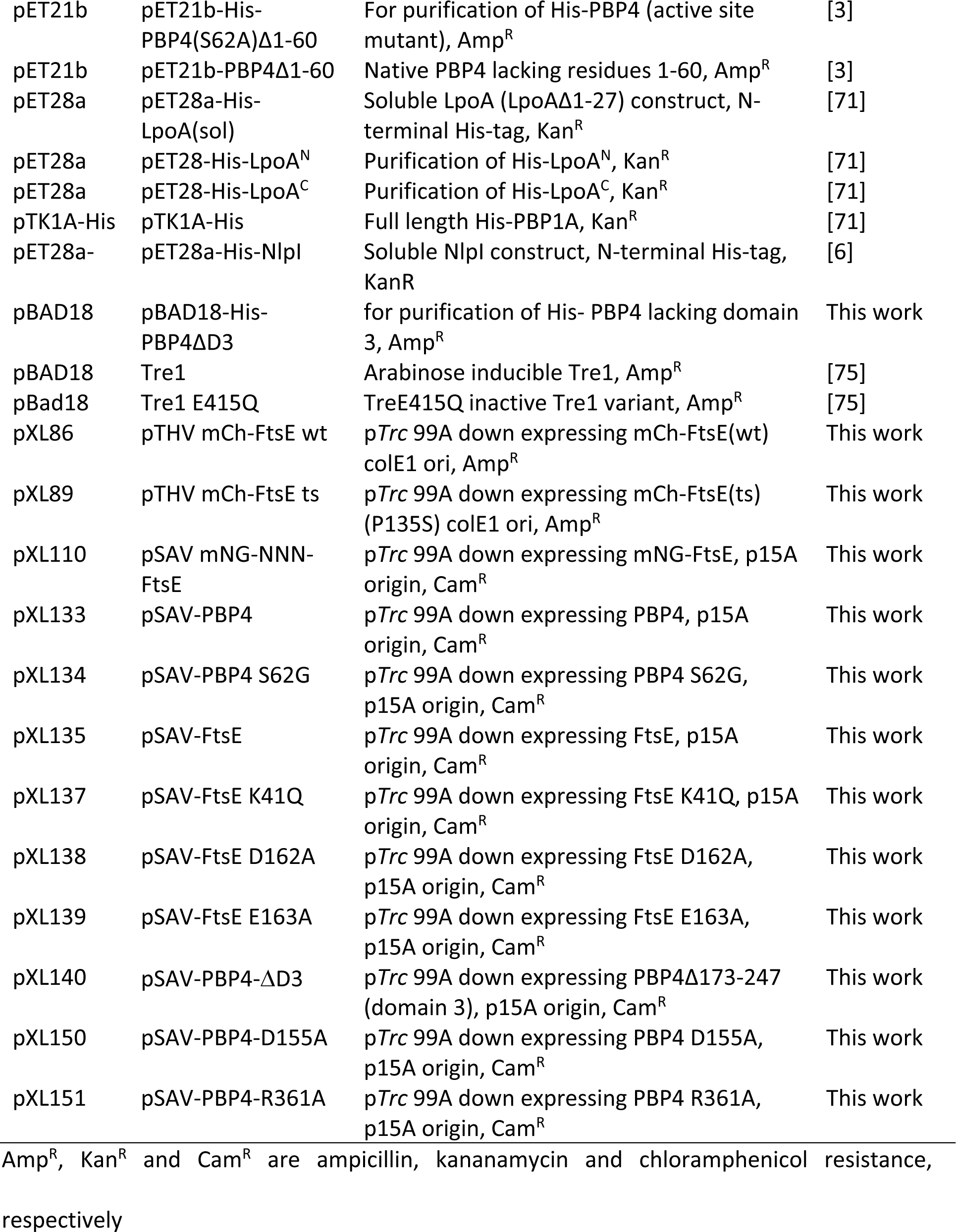
Strain and plasmids used in this study.

### Plasmid construction

The *ftsE* gene was amplified from the chromosome of LMC500[86] using the forward primer priXL187 and the reverse primer priXL188, and sequenced to characterize the mutation. The wild-type *ftsE* gene was cloned into pSAV057, p15A origin and Cam^R^[85] to produce the plasmid pXL135. *FtsE* was expressed under the control of the p*Tcr*99A down promoter. QuickChange site directed mutagenesis (Agilent technologies, Santa Clara, CA) and Gibson assembly[87] approaches were applied afterwards to construct the inactive *ftsE* mutant plasmids pXL137, pXL138 and pXL139, using the primers listed in Table 2.

**TABLE 2.**
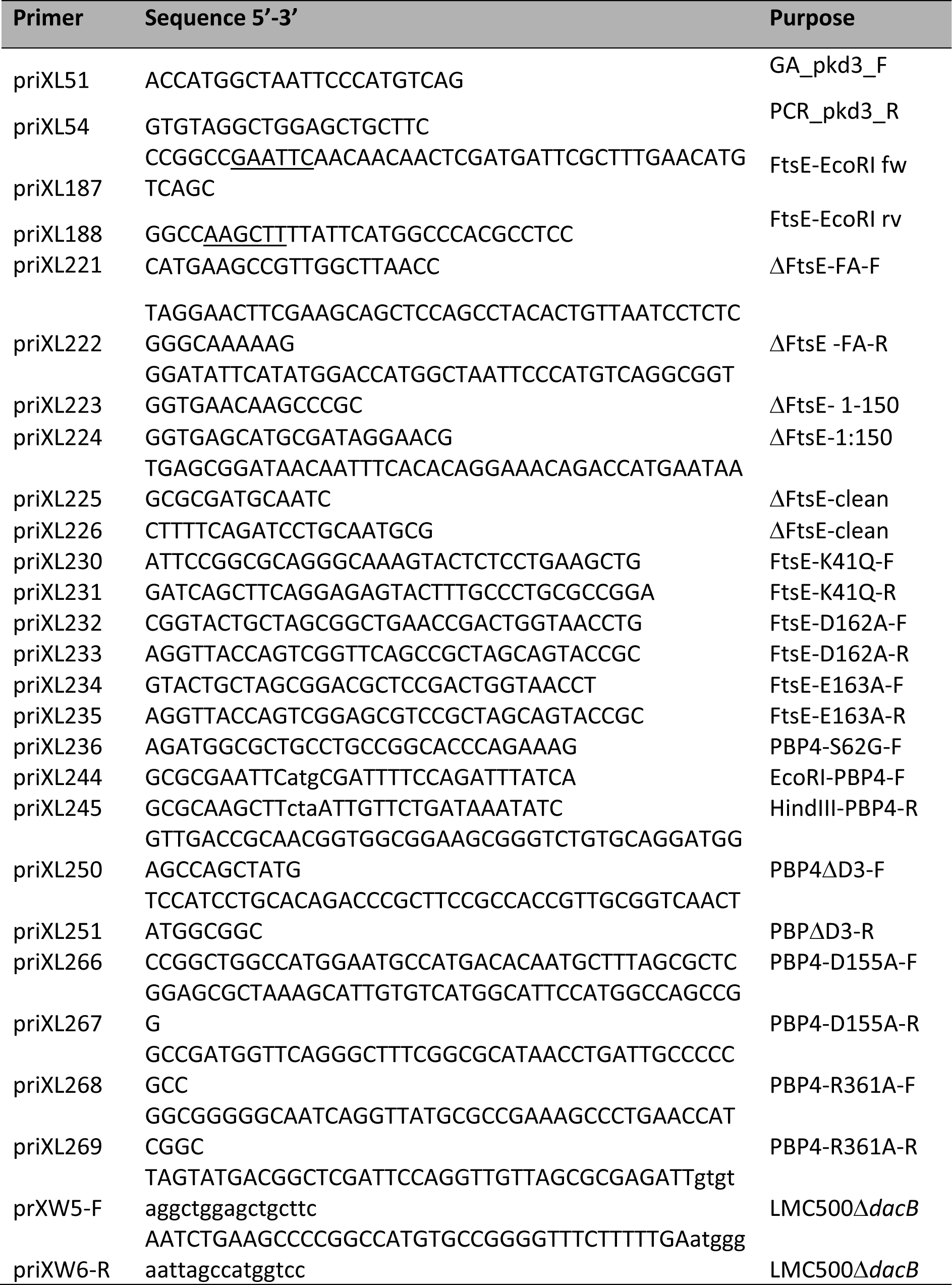
Primers used in this study.

To construct the PBP4 expression plasmids, the wild-type *dacB* gene was firstly amplified from the *E. coli MG1655* genome using primers priXL244 and priXL245, and subsequently cloned into plasmid pSAV057 and pSAV057-dsba^ss^-mCherry-PBP5 (pNM010[20]) with *EcoRI* and *HindIII*, to generate the PBP4 plasmids without and with mCherry fusion. Similarly, QuickChange site directed mutagenesis and Gibson assembly approaches were applied afterwards to construct the inactive *dacB* mutant plasmids from these two wild-type plasmids, using the primers listed in Table 2. Tre1 and Tre1E415Q were expressed from the arabinose inducible promoter of pBAD18 as described[75].

### Immunolabeling

After reaching steady state for minimal glucose grown cells or the desired OD for rich medium grown cells, the cells were fixed for 15 min by addition of a mixture of formaldehyde (f. c. 2.8%) and glutaraldehyde (f. c. 0.04%) to the cultures in the shaking water bath and immunolabeled as described[88] with Rabbit polyclonal antibodies against PBP4, NlpI[3] preabsorbed against *ΔdacB* or *ΔnlpI*, strains, respectively, or against FtsZ or FtsN[48]. As secondary antibody, donkey anti-rabbit conjugated to Cy3 or to Alexa488 (Jackson Immunochemistry, USA) diluted 1:300 in blocking buffer (0.5% (wt/vol) blocking reagents (Boehringer, Mannheim, Germany) in PBS) was used, and the samples were incubated for 30 minutes at 37°C. For immunolocalization, cells were immobilized on 1% agarose in PBS slabs coated object glasses as described[76] and photographed with an Orca Flash 4.0 (Hamamatsu, Japan) CCD camera mounted on an Olympus BX-60 (Japan) fluorescence microscope through a 100x/*N.A*. 1.35 oil objective. Images were taken using the program ImageJ with MicroManager (https://www.micro-manager.org).

SIM images were obtained with a Nikon Ti Eclipse microscope (Japan) and captured using a Hamamatsu Orca-Flash 4.0 LT camera. Phase contrast images were acquired with a Plan APO 100x/1.45 Ph3 oil objective. SIM images were obtained with a SR APO TIRF 100x/1.49 oil objective, using 3D-SIM illumination with a 561nm laser, and were reconstructed with Nikon-SIM software using the values 0.20-0.25-0.20 for the parameters Illumination Modulation Contrast (IMC), High Resolution Noise suppression (HNS) and Out of focus Blur Suppression (OBS), respectively.

### Image analysis

Phase contrast and fluorescence images were combined into hyperstacks using ImageJ (http://imagej.nih.gov/ij/) and these were linked to the project file of Coli-Inspector running in combination with the plugin ObjectJ (https://sils.fnwi.uva.nl/bcb/objectj/). The images were scaled to 15.28 pixel per μm. The fluorescence background has been subtracted using the modal values from the fluorescence images before analysis. Slight misalignment of fluorescence with respect to the cell contours as found in phase contrast was corrected using Fast-Fourier techniques as described[80]. Data analysis was performed as described[80]. In brief, midcell was defined as the central part of the cell comprising 0.8 µm of the axis. From either cell part, midcell and remaining cell, the volume, the integrated fluorescence, and thus the concentration of fluorophores was calculated. The difference of the two concentrations is multiplied with the volume of midcell. It yields FCPlus (surplus of fluorescence) at midcell. For age calculation, all cell lengths are sorted in ascending order. Then the equation

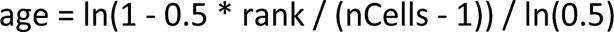

is used, where *rank* is a cell’s index in the sorted array, *nCells* is the total number of cells, and *age* is the cell’s age expressed in the range 0. 1.

### Ni^2+^-NTA pulldown assays

Nickel-Nitrilotriacetic acid (Ni-NTA) beads (100μl) were pre-equilibrated with dH_2_O and binding buffer (10 mM HEPES/NaOH, 10 mM MgCl_2_, 150 mM NaCl, 0.05% Triton X-100, pH 7.5) by centrifugation at 4000 *× g*, 4 min at 4°C, and then incubated with proteins of interest. Equimolar amounts (1-2 μM) of His6-tagged and untagged proteins were incubated alone (control) and in combination, for 10 min at 4°C in a 200 μl reaction volume, containing 10 mM binding buffer. An aliquot of this mixture is taken as an ‘applied’ sample. The protein samples were then added to pre-equilibrated Ni-NTA beads and incubated overnight on a spinning plate at 4°C with 1.3 ml of binding buffer.

Beads were centrifuged at 4000 *× g*, 4 min, 4°C and washed a further 4-6 times with 1 ml of washing buffer (binding buffer with 30 mM Imidazole) before re-suspending in 250 μl washing buffer and transferring beads to Proteus spin columns (Generon), centrifuging as described above. Bound proteins were eluted by the addition of 50 μl of SDS-PAGE loading buffer and boiling at 100°C for 5 min. Spin columns were centrifuged a final time at 1500 *× g*, 5 min, RT, to collect ‘eluted’ protein before separation by SDS-PAGE. Eluted samples were run alongside applied samples for comparison.

### Microscale thermophoresis

Microscale thermophoresis (MST) is an immobilisation-free method which allows the detection of biomolecular interactions in solution. This technique is based on the specific directed movement of a protein along a heat gradient (thermophoresis), an observation first reported by Carl Ludwig in 1856[89]. The thermophoresis of a protein changes upon ligand binding due to one or more changes to size, charge and/or hydration shell. In MST, a localized heat gradient is initiated by an IR-laser (wavelength 1470 nm). A protein of interest is fluorescently labelled and the change in its thermophoretic mobility, in the presence of an unlabelled ligand, is measured and expressed as a change in the normalised fluorescence (FNorm).

Proteins of interest (10-20 μM) were fluorescently labelled with an amine reactive dye (NT-647-N-hydroxysuccinimide (NHS)), cysteine reactive dye (NT-647 maleimide) or a histidine reactive dye (NT-647-Tris-NTA), according to manufacturer’s instructions (Nanotemper). Fluorescently labelled proteins were diluted to an appropriate concentration and a fixed concentration was titrated against a two-fold serial dilution of unlabelled ligand, across 16 samples, in MST running buffer (25 mM HEPES/NaOH, 150 mM NaCl, 0.05% Triton X-100, pH 7.5).

MST measurements were carried out as described in[90], using standard or premium capillaries on a Monolith NT.115^TM^ MST machine (Nanotemper). Binding curves and Kinetic parameters were plotted and estimated using manufacturer provided software (NT Analysis 1.5.41 and MO. Affinity Analysis (x64)). Capillary scans were carried out prior to all measurements to check for consistent fluorescence counts, confirming that any subsequent change in fluorescence was due to ligand binding and not due to inaccurate pipetting or adsorption and dilution effects.

### SDS-denaturation (SD) test

In instances where capillary scans showed a ligand concentration dependant change in raw fluorescence, we investigated whether this was a property of the binding interaction by carrying out an SD-test.

Samples (10 μL) with the highest and lowest concentration of unlabelled ligand were centrifuged 10 000 × *g,* 5 min, RT and mixed 1:1 volume ratio with SD-test buffer (40 mM DTT, 4% SDS). Mixtures were boiled at 100 °C for 10 min, to abolish ligand binding, before being spun down and subjected to another capillary scan. If the fluorescence between samples that contained the highest and lowest concentration of ligand, after SDS treatment, were now back to ± 10% of each other, the initial observations were a property of ligand binding, and a binding curve was plotted from the raw fluorescence data. If the ligand concentration dependent change in fluorescence was still observed, then this indicated that the fluorescently labelled protein was aggregated, and assay and buffer conditions were optimised accordingly

### *In vivo* cross-linking / co-immunoprecipitation assays

Method is described in, and adapted from[91]. An overnight culture of *E. coli* BW25113 cells and an appropriate mutant strain was used to inoculate 150 ml of Lennox LB (Fisher Scientific) and was cultivated to an OD_578_ of 0.5-0.6 at 37°C before harvesting by centrifugation (4500 × g, 4°C, 25 min). Cells were resuspended in 6 ml of CL buffer 1 (50 mM NaH_2_PO_4_, 20% sucrose, pH 7.4). The amine reactive cross-linker, DTSSP (3,3’-dithiobis (sulfosuccinimidylpropionate) (ThermoFisher), was freshly dissolved (20 mg/ml in dH_2_O) and added to the isolated cell suspension and incubated at 4°C with agitation for 1 h. Cross-linked cells were then harvested by centrifugation (4500 × g, 4°C, 25 min) and resuspended in 6 ml CL buffer 2 (100 mM Tris/HCl, 10 mM MgCl_2_, 1 M NaCl, pH 7.5). DNase, protease inhibitor cocktail and phenylmethylsulfonly fluoride were added prior to sonication at low levels before ultracentrifugation of the lysate (140,000 × g, 4°C, 1 h). The membrane pellet was resuspended in 2.5 ml of CL buffer 3 (25 mM Tris/HCl, 10 mM MgCl_2_, 1 M NaCl, 1% Triton X-100, 20% glycerol, pH 7.5) and the solubilised membrane extracted o/n with stirring at 4°C.

Samples were ultracentrifuged (140,000 *× g*, 4°C, 1 h) to remove debris, before removing 2 × 1.2 ml of each supernatant to be subsequently diluted with 0.6 ml of CL buffer 4 (75 mM Tris/HCl, 10 mM MgCl_2_, 1 M NaCl, pH 7.5). One sample was incubated with an optimised concentration of specific antibody with the other used as a negative control. Both samples were incubated at 4°C with agitation for 5 h. For the isolation of antibodies, and thus cross-linked interaction partners, 100 µl of protein G-coupled agarose bead resin (Roche) were washed (2 × CL buffer 4, 2 × CL wash buffer [2:1 CL buffer 3 and CL buffer 4]) and added to each sample, and incubated o/n at 4°C with agitation.

Samples were centrifuged and the supernatant retained before washing the beads 10 × 1 ml with CL wash buffer. After the final wash, beads were resuspended in 250 µl CL wash buffer and transferred to 2 ml spin dry columns and centrifuged to isolate the beads. These were then resuspended in 50 µl of fresh SDS-loading buffer and boiled to elute bound proteins, and reverse cross-linkage, and were collected by centrifugation (10,000 *× g*, RT, 5 min). Supernatant and elution samples were resolved by SDS-PAGE and transferred to a nitrocellulose membrane by Western blotting to detect for specific interaction partners using purified antibodies. The secondary antibody used here is Trueblot Anti-Rabbit IgG-HRP specific for native antibodies.

### Peptidoglycan digestion assays

To test for hydrolase activity on PG, 10 μl of sacculi isolated from strains of interest (usually *E. coli* strains MC1061, BW25113 or D456) were incubated with 1-10 μM of respective enzymes, 37°C for between 1 h and overnight, as indicated for respective proteins. The standard reaction conditions were 10 mM HEPES/NaOH, 10 mM MgCl_2_, 150 mM NaCl, 0.05% Triton X-100, pH 7.5, in 100 μl reaction volume. Following incubation, samples were boiled at 100°C, 10 min, to terminate reactions before digesting remaining PG overnight at 37°C, with 1 μM cellosyl. The samples were centrifuged at 10,000 × *g* for 5 min, RT, to obtain digested muropeptide products in the supernatant.

To test for activity against soluble muropeptides; first, 100 μl of intact sacculi were incubated overnight at 37°C with 1 μM cellosyl and cellosyl digestion buffer (20 mM NaPO4, pH 4.8). Next, samples were boiled at 100 °C, 10 min, before centrifugation at 10,000 × *g* for 5 min, RT, to obtain the soluble muropeptides in the supernatant[92]. Ten μl of the supernatant was then used as muropeptide substrate for incubating with enzymes of interest, for the appropriate incubation time, at 37°C. Reactions were then terminated by boiling at 100°C for 10 min.

After digestion of respective substrates, products were reduced with NaBH4, adjusted to pH 4-5 and separated for analysis by reversed-phase HPLC as described below in HPLC analysis.

### Reduction of muropeptides with sodium borohydride

PG digestion samples were transferred to 2 ml vials following centrifugation. Muropeptides were reduced in a 1:1 volume ratio of sodium borohydride buffer (0.5 M sodium borate, pH 9.0) and a small spatula of sodium borohydride pellets, centrifuging at 3000 × *g,* 30 min, RT as described in[92]. The samples were adjusted to pH 4-5 using HPLC grade phosphoric acid before separation and analysis by reversed-phase HPLC.

### Reversed-phase HPLC analysis of muropeptides

Following the protocol of[92], reduced muropeptides were separated for analysis on reversed-phase HPLC systems with a Prontosil 120-3-C18-AQ 3 μm reversed-phase HPLC column (Bischoff). A linear gradient of solvent A (50 mM sodium phosphate, pH 4.31 supplemented with 0.2% NaN_3_) to 100 % solvent B (75 mM sodium phosphate, pH 4.95, 15% methanol) over 90 or 180 min was used to separate muropeptides at 52°C. Unlabelled muropeptides were detected at UV absorbance 205 nm. In assays where ^14^C-radiolabelled muropeptides were used, detection was achieved by flowing scintillation cocktail along with standard buffers to give a radioisotope scintillation count (radioactivity CPM). Muropeptide profiles were recorded and analysed using Laura V4.2.11.129 (LabLogic Systems Ltd.).

### Spectrophotometric D-alanine release assay (DD-carboxypeptidase assay)

This protocol was adapted from[62]. The carboxypeptidase (CPase) activity of PG hydrolases results in the release of the terminal D-Ala residue from the pentapeptide stem of PG precursors. Using UDP-Mur*N*Ac pentapeptide as a substrate, and in this case PBP4, it was possible to spectrophotometrically measure the release of D-Ala.

Each reaction sample consisted of 200 μl of CPase buffer (50 mM HEPES/NaOH, 10 mM MgCl_2_, pH 7.6), 3 units of D-amino acid oxidase (Sigma), 6 units of horseradish peroxidase (HRP) (Sigma), Amplex Red (Sigma), and an optimised concentration of protein.

All constituents of the reaction were added and mixed directly in a quartz cuvette (Hellma, 10 mM light path, 15 mM centre), before the addition, and brief mixing by pipette, of purified UDP-Mur*N*Ac pentapeptide (BACWAN, Warwick University) to begin the reaction. The released D-Ala residues from the CPase activity of PBP4 are oxidatively deaminated by the action of D-amino acid oxidase to produce pyruvate and hydrogen peroxide (H_2_O_2_). The released H_2_O_2_ is reduced to H_2_O by HRP using Amplex Red as an electron donor. Oxidised Amplex Red produces resorufin, which has an intense pink colour and the production of which was measured spectrophotometrically using a Cary 100 Bio UV-visible spectrophotometer (wavelength 555 nm). The change in absorption over 10 min was measured and analysed using the complementing software.

### Circular dichroism

Proteins were dialysed overnight against 10 mM NaPO4, pH 7.5 and concentrated/diluted to 0.4 mg/ml. CD measurements were taken using a Jasco J-810 spectropolarimeter using a wavelength range of 180-250 nm. The average of 10 runs was taken for each protein with a buffer control subtraction. For a direct comparison, correcting for the differing amino acid sequences, the collected data were converted to molecular CD and plotted against wavelength (nm). The resulting CD spectra are compared in Fig. S4 and show that PBP4 lacking domain 3 is folded, consisting of both α-helices (∼190, 208 and 222 nm) and β-sheets (∼210 nm).

### Analytical ultracentrifugation

Purified PBP4 and PBP4ΔD3 were dialyzed over night against 25 mM HEPES/NaOH, 150 mM NaCl, pH 7.5, in preparation for AUC experiments. AUC sedimentation velocity (SV) experiments were carried out in a Beckman Coulter (Palo Alto, CA, USA) ProteomeLab XL-I analytical ultracentrifuge using absorbance at 280 nm and interference detection. The AUC runs were executed at the rotation speed of 45,000 rpm and the temperature of 20°C using an 8-hole AnTi50 rotor and double-sector aluminium-Epon centerpieces. The sample volume was 400 µl and the sample concentrations ranged between approximately 0.25 and 1.4 mg/ml. The partial specific volumes (v̄) of the proteins were calculated from their amino acid sequence, using the program SEDNTERP[93]. Sedimentation velocity boundaries were analyzed using the size-distribution c(s) model implemented in the program SEDFIT http://www.analyticalultracentrifugation.com[94]. The experimental values of the sedimentation coefficient were converted to the standard conditions (s20,w), which is the value of sedimentation coefficient in water at 20°C. The size-distribution peaks were integrated to obtain the weight-averaged values for sedimentation coefficient and molecular mass.

The atomic coordinates from the published high-resolution structure of[60] (pdb accession code 2EX2) were used to calculate the sedimentation coefficient values for the monomeric and dimeric forms of PBP4 /PBP4ΔD3 using the program SoMo[95]. PBP4/PBP4ΔD3 crystallographic dimer was built using program PyMol (https://pymol.org/2/).

### PG binding assay

To assay for interactions of respective proteins with purified PG, we used a PG binding protocol as described in[96]. Briefly, ∼100 μg purified PG was pelleted by centrifugation (10,000 × *g,* 10 min, 4°C) and resuspended in binding buffer (10 mM Tris/Maleate, 10 mM MgCl_2_, 50 mM NaCl, pH 6.8). PG was incubated with 10 μg of desired protein before incubating on ice for 30 min and centrifuging at 10,000 × *g,* 10 min, 4°C. An aliquot of the supernatant (S) was taken for SDS-PAGE. The pelleted PG was washed with 200 μl binding buffer by centrifugation as before and the wash supernatant was taken as sample W for SDS-PAGE. The pelleted PG was then resuspended in 100 μl of 2% SDS and incubated for 1 h on a stirring plate, 4°C. Mixtures were centrifuged a final time at 10,000 × *g,* 10 min, 4°C, with the supernatant, now containing any initially bound protein, taken as the pellet sample (P). A negative control sample with no PG was assayed in parallel to determine that binding of protein was specific. Samples, along with corresponding controls, were analysed by SDS-PAGE and visualised by Coomassie staining. Presence of protein band in pellet sample (P) indicated binding to PG, whilst presence in supernatant sample (S) indicated no binding.

### Bocillin binding assay

A fluorescent-bocillin binding assay was used to determine whether a PBP was purified with a correctly folded active site. Bocillin FL is a commercially available dye-β-lactam conjugate, synthesised from penicillin V and BODIPY (Molecular Probes, In., Eugene, Oreg.). Ten μg of purified PBP was incubated for 10 min at 37°C, with 20 ng of Bocillin FL in a 50 μl volume with 10 mM HEPES/NaOH, 10 mM MgCl_2_, pH 7.5. A negative control sample was pre-incubated with 1 mM ampicillin for 30 min, 37°C, to block the PBP active site, prior to incubation with Bocillin FL. After incubation, samples were boiled at 100°C for 10 min with 40 μl of SDS-PAGE loading buffer and resolved by SDS-PAGE. Fluorescent signal was observed using Typhoon Fluorescence-imager (Excitation; 488 nm, emission; 520 nm, BP20 PMT 400-600 V), followed by visualisation of gels by Coomassie staining. Proteins incubated with ampicillin should have no detectable binding of Bocillin FL. For whole cells, a culture was grown in 5 mL TY and induced with 10 μM IPTG for 3 mass doublings to an OD_600_ of 0.5 and the cells were pelleted and washed once with PBS (8 g NaCl, 1.8 g Na_2_HPO_4_, 0.24 g KH_2_PO_4_ per liter, adjusted to pH 7.4) at 4°C and resuspended in 100 μl PBS with 5μg/ml Bocilin FL and incubated for 20 min at RT. To remove the excess Bocillin FL, the samples were pelleted and resuspended in 100 μl PBS with cOmplete™ Mini Protease Inhibitor Cocktail (Sigma) and 2 U DNAse I (NEB). Then sonicated on ice for 10 sec and boiled 15’ at 99°C after adding 25 μL of SDS-PAGE 5x loading buffer and resolved by SDS-PAGE. Fluorescent signal was observed using a home-made imager for Midori green, followed by visualisation of gels by Coomassie staining.

### *In vitro* PG synthesis assay with radiolabelled lipid II

The assay was published by Bertsche et al.,[97]. Lipid II (1.2 nmol, 11,000 dpm) was vacuum dried and dissolved in 5 µl of methanol. The reaction was performed in a total volume of 50 µl in 25 mM Tris/HCl, 10 mM MgCl_2_, 100 mM NaCl, 0.05% Triton X-100, pH 7.5 (buffer B), with 0.5 µM PBP1A, 1 µM LpoA, 1 µM PBP4(S62A), 2 µM NlpI. The reaction was performed for 1 h at 30°C. The pH of the samples was adjusted to 4.8 prior to boiling for 5 min followed by a digestion with 50 g/ml cellosyl for 1 h at 37°C and boiling for 5 min.

## ACKNOWLEDGEMENTS

We thank David Roper and Smita Chauhan (University of Warwick) for help with the carboxypeptidase assay, Helen Waller (Newcastle University) for help with circular dichroism methodology, Lisa Atkinson and Daniela Vollmer (Newcastle University) for preparation of PG sacculi, and Thomas van Wingen for with help the Tre1 assay. We thank See-Yeun Ting for the plasmids expressing Tre1 and Tre1E415Q (University of Washington School of Medicine, Seattle, USA). The work was supported by funding from the Wellcome Trust (101824/Z/13/Z), to WV). X.L. was supported by the China Scholarship Council fellowship (File No.201406220123). J.V., A.L., H.C.L.Y, X.L., X.L., A.S.S., and M.B did most experiments. A.T., M.B., V.W., and T.dB. designed most experiments. M.B, W.V and T.dB analyzed the data and wrote the manuscript. All authors critically read the manuscript and provided feedback.

## Supplementary figures

**FIG S1.**
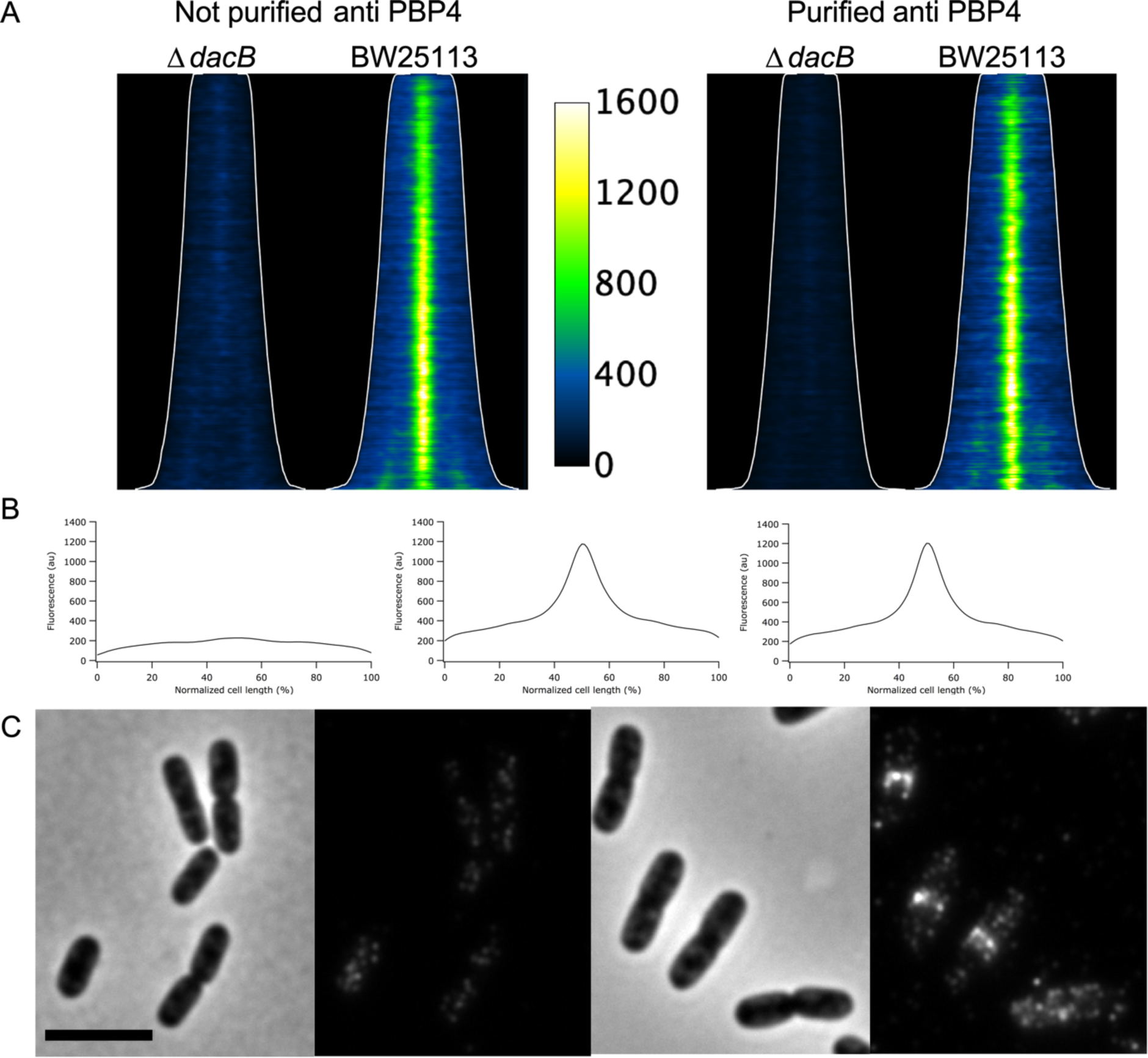
Pre-adsorption of antibodies against Δ*dacB* BW25113 strain results in PBP4 specific antibodies. (A) Demograph with cells sorted according to length of PBP4 fluorescence of the BW25113Δ*dacB* (PBP4) strain, BW245113 (wt) strain before purification of the antibody and the BW25113Δ*dacB* strain and BW245113 strain after purification of the antibody. (B) Absolute fluorescence average profiles of cells from the BW25113 Δ*dacB* strain and the BW245113 strain before purification of the antibody and from the BW245113 strain after purification of the antibody (C) Phase contrast and corresponding fluorescence image of anti PBP4 immunolabeled BW25113Δ*dacB* (left) and wild-type cells using the supernatant of BW25113Δ*dacB* pre-adsorbed antibodies. The scale bar equals 5 µm.

**FIG S2.**
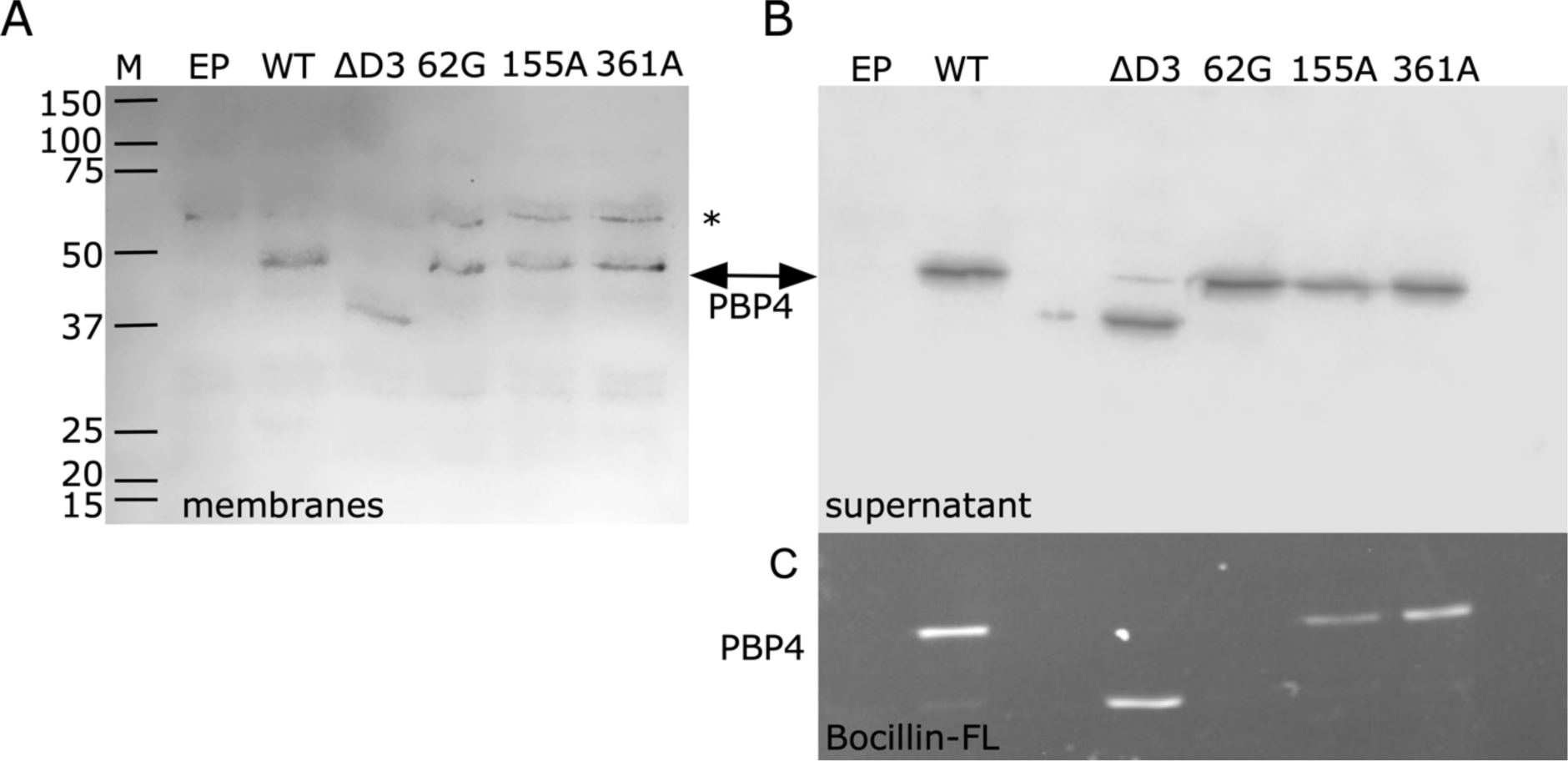
All PBP4 mutants are equally well expressed and only S62G is not able to bind Bocillin-FL. (A) An immunoblot of membranes of PBP4 wild-type and variants. *This extra band is due to non-specificity of the primary antibody, which was for the purpose of immunoblotting not affinity purified. (B) Immunoblot of supernatant after pelleting the membrane of PBP4 wild-type and variants. (C) The corresponding gel where binding of the fluorescent β-lactam Bocillin-FL is visible for all mutants apart from S62G. EP is empty plasmid, WT is wild-type. The other samples are PBP4 variants.

**FIG S3.**
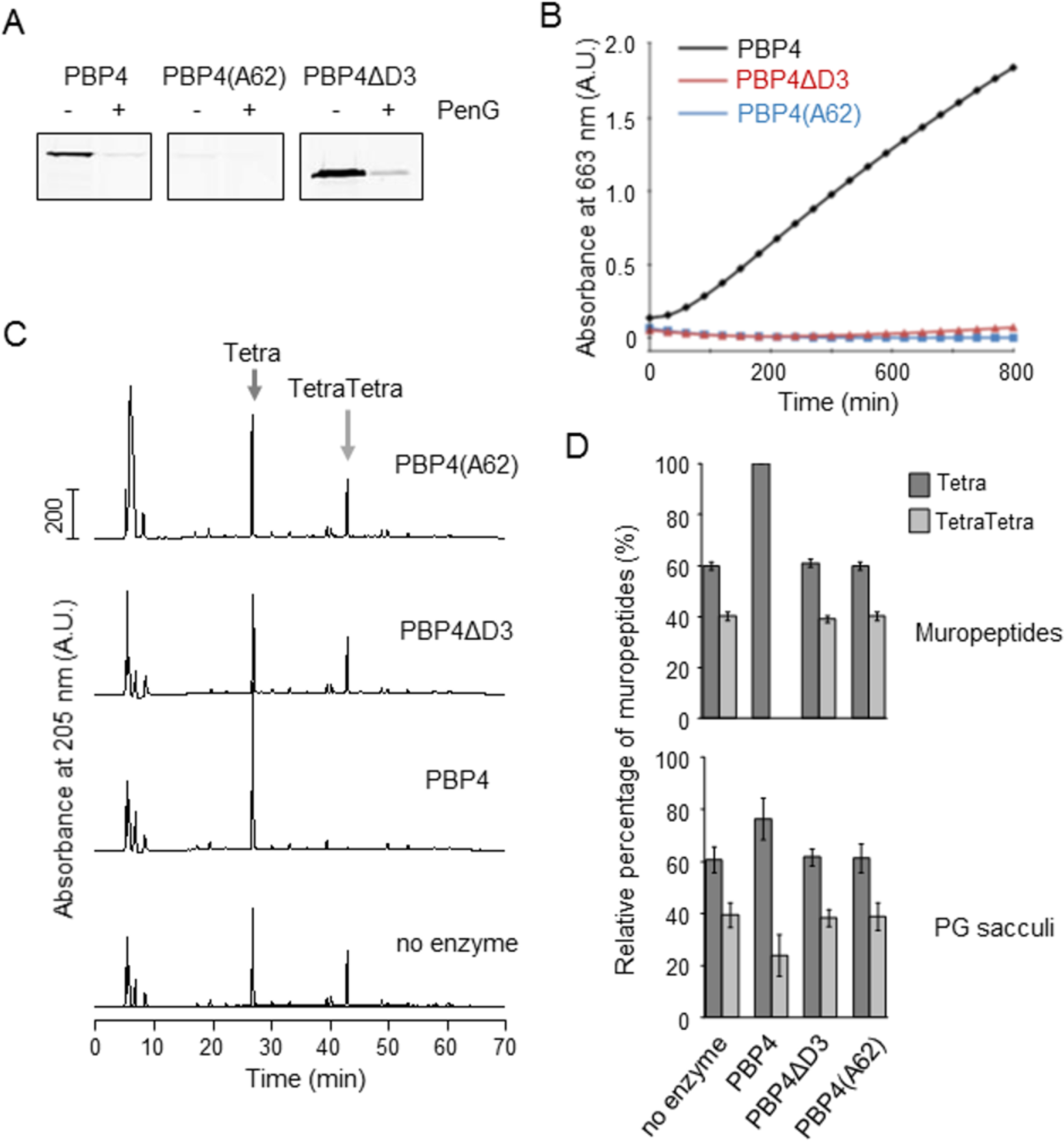
PBP4ΔD3 binds beta-lactam but is inactive against PG substrates. (A) PBP4 versions were incubated with the fluorescent β-lactam Bocillin FL with or without pre-incubation with Penicillin G (PenG), followed by SDS-PAGE analysis and detection of covalent Bocillin-PBP4 adducts by fluorescence scanner. PBP4 and PBPP4ΔD3, but not catalytically inactive PBP4(S62A), bound Bocillin FL. (B) Only wild-type PBP4, but not PBPP4ΔD3 or PBP4(S62A), was active in a DD-carboxypeptidase assay. (C) PBP4 versions were incubated with muropeptides from BW25113 prior to their separation by high-performance liquid chromatography (HPLC). A control sample contained no enzyme. PBP4 DD-endopeptidase activity is demonstrated by the reduction in the dimer (TetraTetra) substrate peak and increase in the monomer (Tetra) product peak. PBPP4ΔD3 and PBP4(S62G) were inactive. (D) Quantification of the Tetra and TetraTetra peaks shown in panel C (top) and quantification of a similar analysis with PG sacculi (incubated with PBP4 versions or no enzyme), followed by generation of muropeptides and HPLC analysis (bottom).

**FIG S4.**
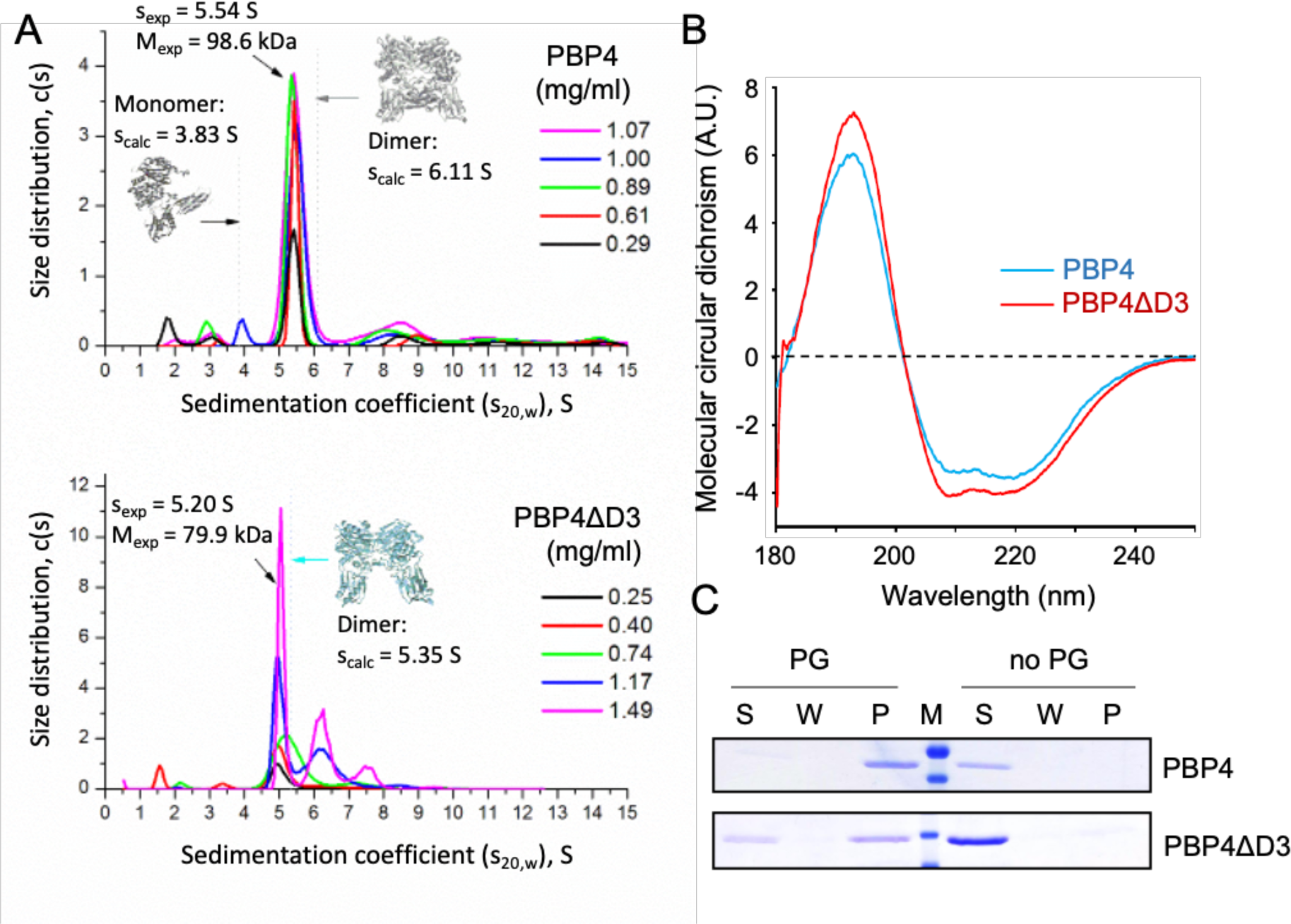
The absence of domain 3 does not affect the dimerization, secondary structure or PG binding of PBP4. (A) Analytical ultracentrifugation sedimentation velocity experiment of PBP4 and PBP4ΔD3 shows that both proteins are mainly dimers. The determined sedimentation coefficients (sexp) for the dimers fit well with the theoretical values (scalc) that were calculated from atomic coordinates of both protein dimers (pdb accession code for monomer is 2EX2). (B) PBP4 and PBP4ΔD3 show similar far UV circular dichroism spectra. (C) PBP4 and PBP4ΔD3 co-sediment with PG sacculi from BW25113 cells, but did not sediment in control samples without sacculi, demonstrating binding to PG. S, supernatant; W, wash fraction; P, pellet fraction.

**FIG S5.**
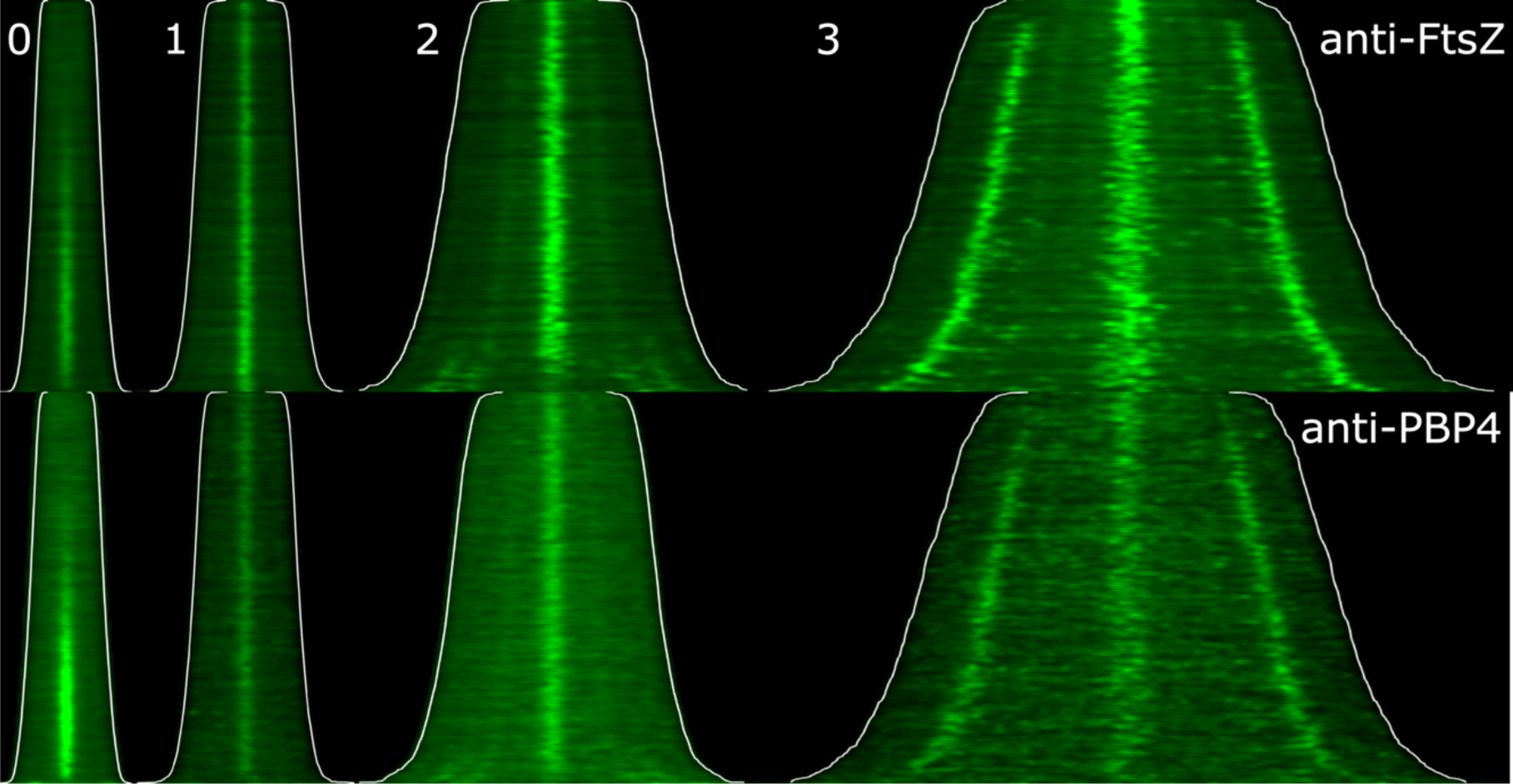
PBP4 localizes at site of inactive divisomes. MC4100 cells were grown to steady state in minimal glucose medium at 28°C and split in two parts. One part was 1:4 diluted in prewarmed medium without aztreonam and the other part was 1:4 diluted in medium with 10 µg/ml aztreonam. The cells continued to grow for 0, 1, 2 or 3 mass doublings (MD) and were fixed and immunolabeled with antibodies specific for FtsZ or PBP4. Demographs with identical brightness and contrast of the FtsZ or the PBP4 fluorescence of cells grown in the absence or presence of aztreonam sorted according to length. The white line shows the borders of the cells. Number of analyzed cells per demograph for FtsZ and PBP4 were, 5084 and 3448 (0), 1430 and 1858 (1), 857 and1630 (2) and 659 and 595 (3), respectively.

**FIG S6.**
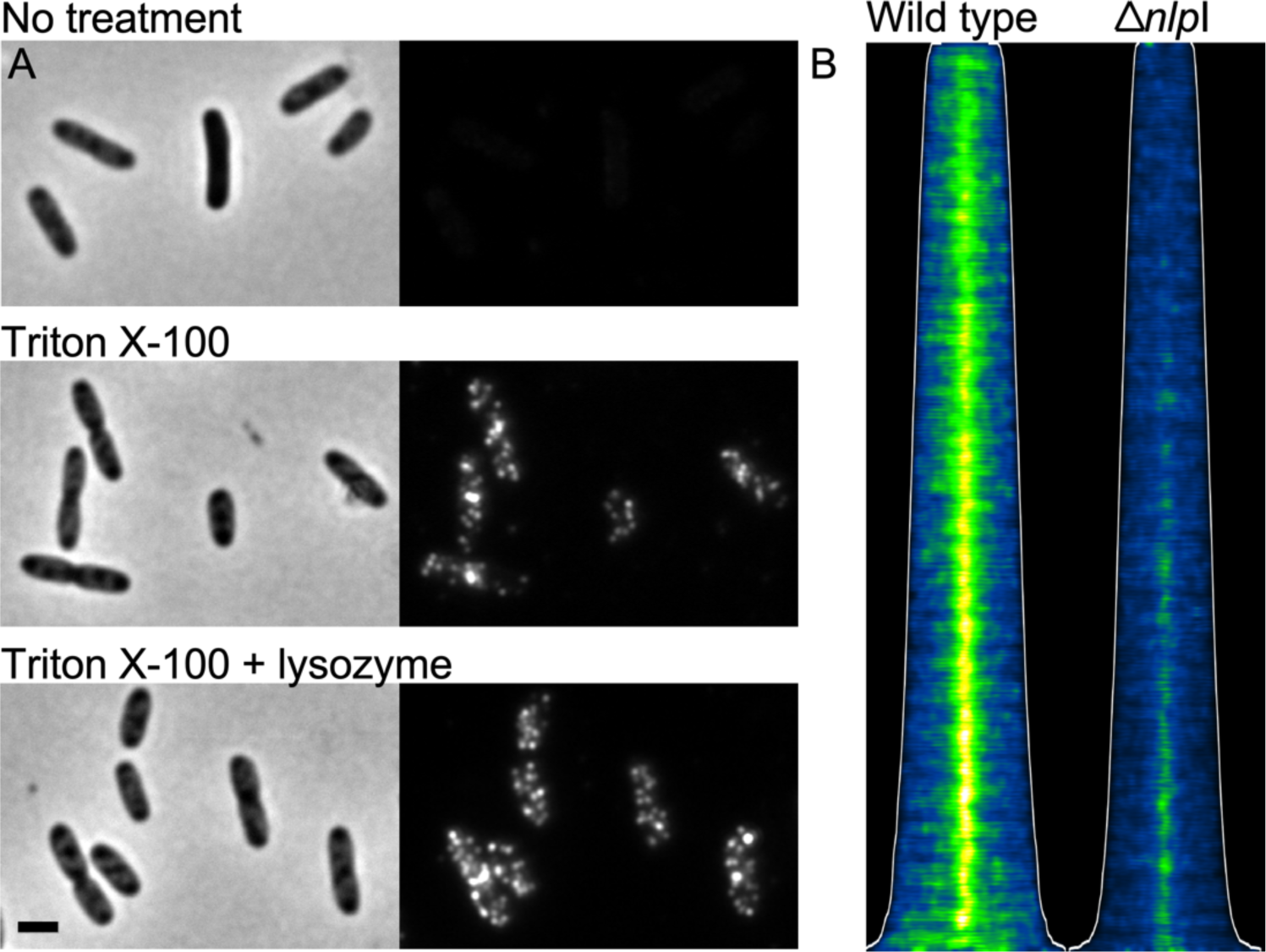
PBP4 localizes between the outer membrane and the peptidoglycan layer in a Δ*nlpI* strain. (A) BW 25113 Δ*nlpI* cells were grown exponentially in TY medium at 37°C and fixed while shaking when at an OD_600_ of 0.3. The cells were harvested and split in three portions of which one was directly immunolabeled with to Δ*dacB* cells pre-adsorbed anti-PBP4, the second was first treated with Triton X-100 and the last was treated with Triton X-100 and lysozyme and then immunolabeled. The scale bar equals 2 μm. (B) Map of PBP4 fluorescence sorted according to cell length of wild-type cells and Δ*nlpI* cells displayed

**FIG S7.**
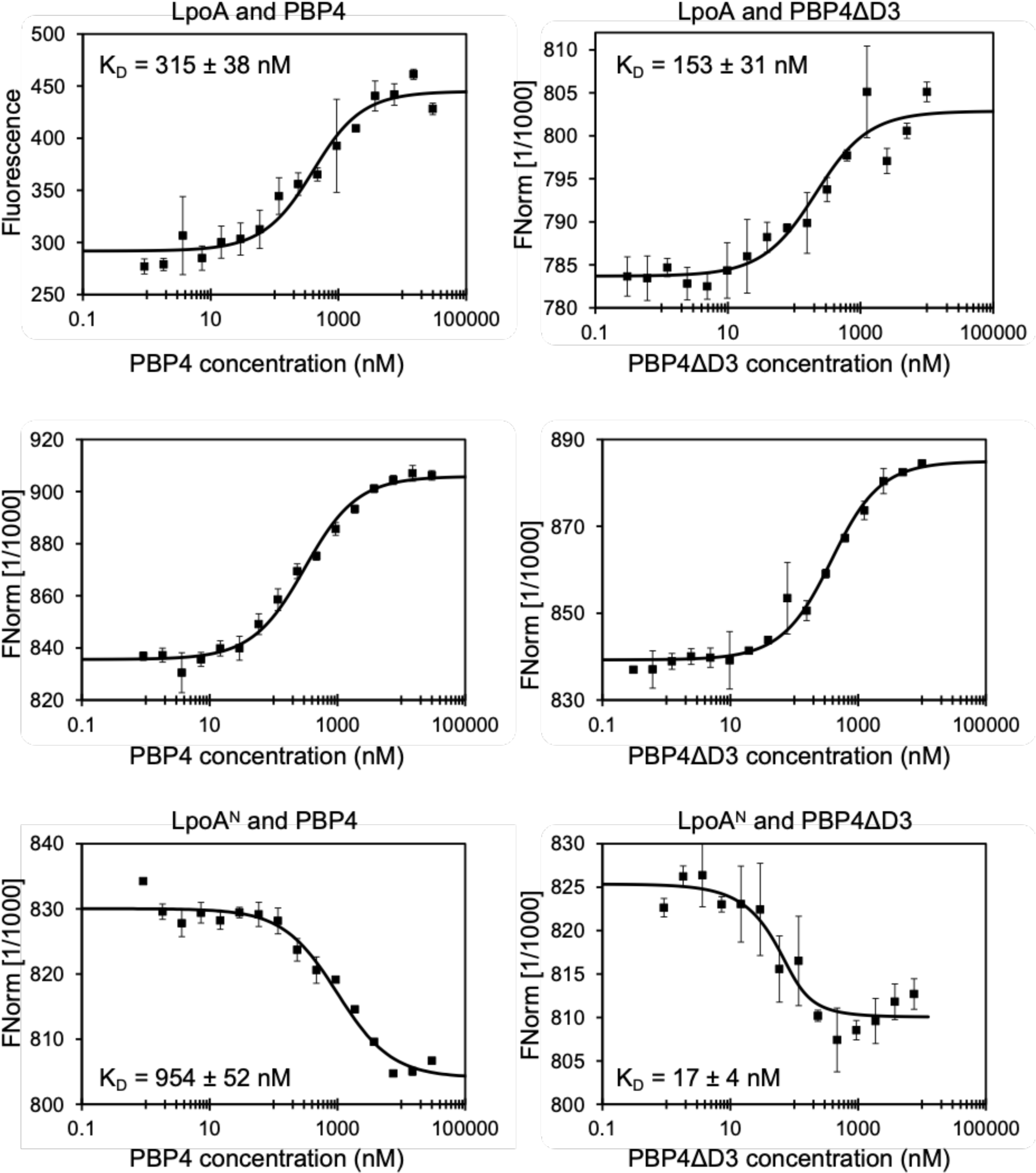
Interactions of PBP4 and PBPP4ΔD3 measured by microscale thermophoresis (MST). MST binding curves for interactions using different fluorescently-labelled proteins (LpoA, LpoA^C^, Lpo^N^, NlpI or PBP1A) titrated against fixed concentrations of PBP4 or PBPP4ΔD3. Apparent K_D_ values are mean ± SD of three independent experiments and summarized in Fig. 4A.

**FiG S8.**
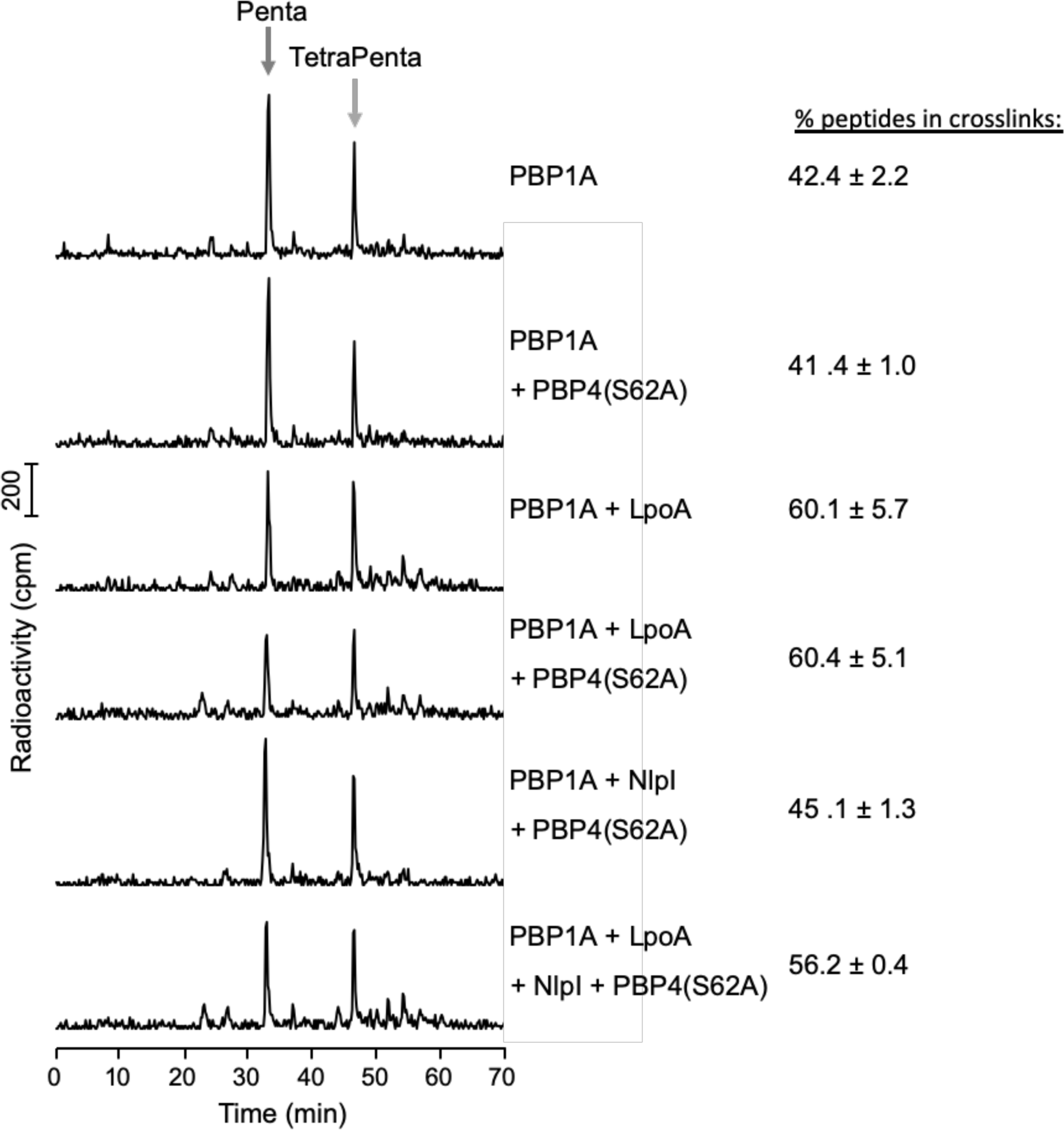
PBP4 or NlpI do not affect the activity of PBP1A or PBP1A/LpoA. Representative HPLC chromatograms of samples from an *in vitro* PG synthesis assay of PBP1A and radiolabelled lipid II, in the presence of proteins indicated. The main PG products upon digestion with the muramidase cellosyl are the disaccharide pentapeptide (Penta) and the bis-disaccharide tetrapentapeptide (TetraPenta). The % peptides present in cross-links is quantified on the right side and presented as mean ± variation of two independent experiments.

**FIG S9.**
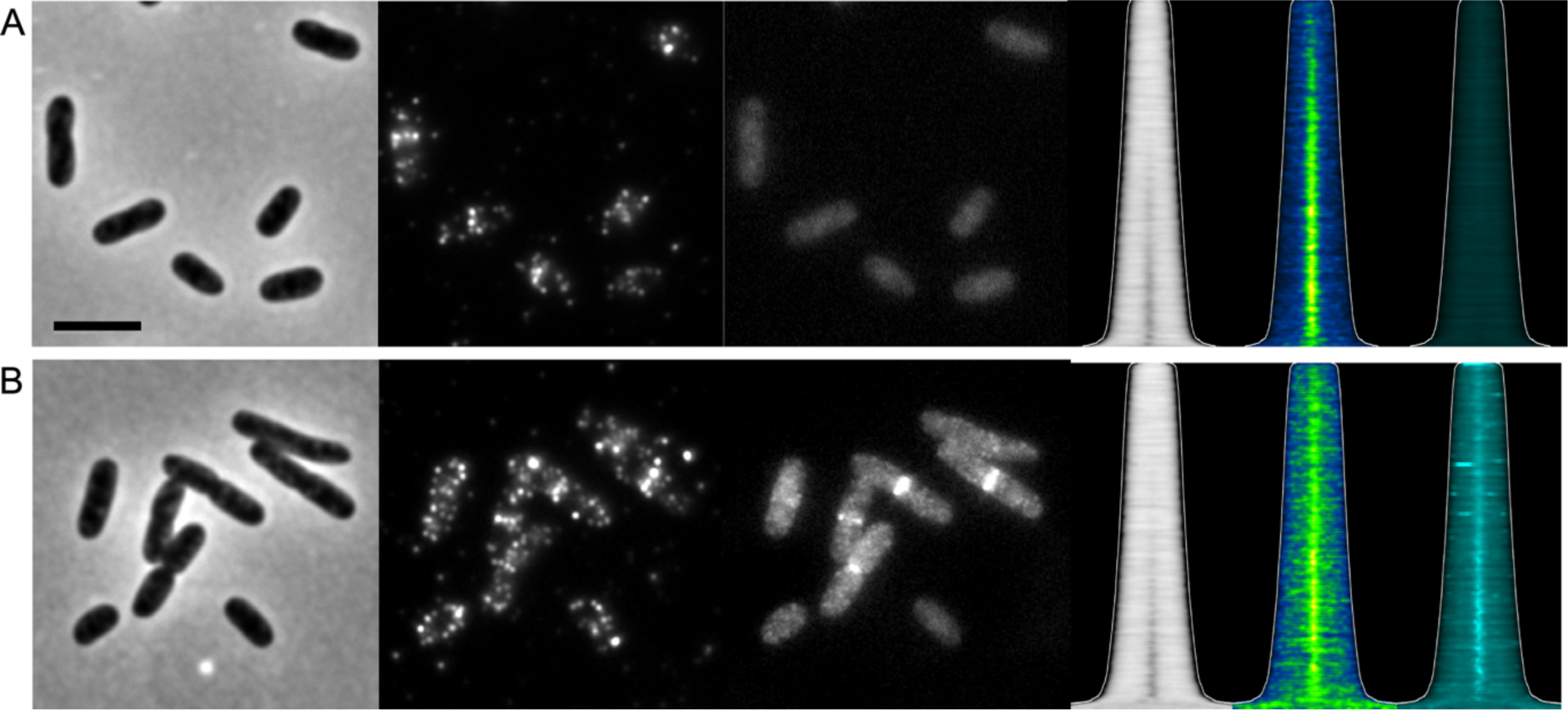
NG-FtsE effectively localizes and recruits PBP4. Strains were grown in LB at 37°C to an OD of 0.3 in the presence of 30 µM IPTG to induce expression of mNG-FtsE then fixed and immunolabeled with antibodies specific for PBP4. (A) LMC500 (1108 cells analyzed). (B) XL36 (LMC500::p*Trc*99down*ftsX*, Δ*ftsE*) with pXL110 mNG-FtsE(wt) (1189 cells analyzed). From left to right, phase contrast image, anti-PBP4 immunolabeling, mNG-FtsE, and the corresponding map of diameter with the fluorescence maps of cells sorted according to length. Scale bar equals 5 µm.

**FIG S10.**
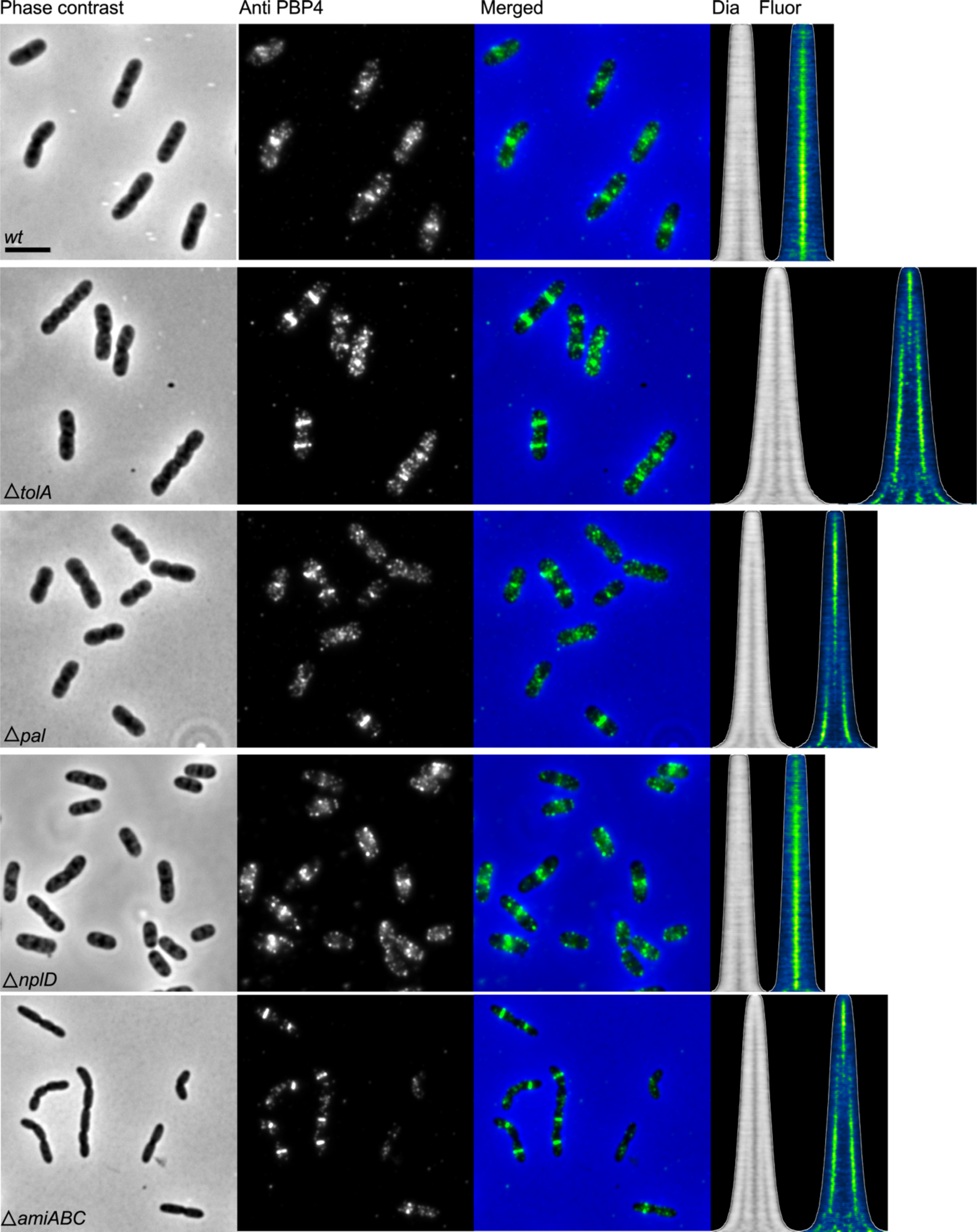
PBP4 localization is not dependent on the presence of TolA, Pal, NlpD or AmiABC. Isogenic strains of the wild-type strain BW25113 were grown in LB at 37°C to a OD_600_ of 0.3, fixed and labeled with specific antibodies against PBP4. From left to right, the phase contrast, corresponding fluorescence image of the PBP4 labeling, and the merged former two images, map of diameters (Dia) and map of fluorescence (Fluor) PBP4 localization where cells are sorted according to their cell length are shown. The number of cells analyzed were 1053 for BW25113 (*wt*), 918 for Δ*tolA*, 1314 for Δ*pal*, 1600 for Δ*nlpD*, and 1410 for Δ*amiABC*. The scale bar equals 5 µm.

**Table S1.** Morphological parameters LMC500 and LMC500Δ*dacB*.

| Parameter | LMC500 | n | LMC500$\Delta dacB$ | n | p |
| --- | --- | --- | --- | --- | --- |
| Length ($\mu m$) | 2.57 ± 0.07 | 8 | 2.58 ± 0.09 | 12 | 0.79 |
| Diameter ($\mu m$) | 0.89 ± 0.09 | 8 | 0.91 ± 0.19 | 12 | 0.73 |
| Constrictions (%) | 16.9 ± 1.8 | 8 | 17.4 ± 1.3 | 12 | 0.53 |

n = number of biological replicates, P from a two tailed T-test to verify whether that the means are significant different.

## Notes

### Competing Interest Statement

The authors have declared no competing interest.

### Summary of Updates

The paper is essentially the same, some data that were not so important have been taken out to reduce the size of the manuscript. The analysis of the divisome timing in the absence of PBP4 has been extened. The conclusions are the same.

